# Myeloid Lineage Enhancers Drive Oncogene Synergy in CEBPA/CSF3R Mutant Acute Myeloid Leukemia

**DOI:** 10.1101/639617

**Authors:** Theodore P. Braun, Mariam Okhovat, Cody Coblentz, Sarah A. Carratt, Amy Foley, Zachary Schonrock, Kimberly Nevonen, Brett Davis, Brianna Garcia, Dorian LaTocha, Benjamin R. Weeder, Michal R. Grzadkowski, Joey C. Estabrook, Hannah G. Manning, Kevin Watanabe-Smith, Jenny L. Smith, Amanda R. Leonti, Rhonda E. Ries, Sophia Jeng, Shannon McWeeney, Cristina Di Genua, Roy Drissen, Claus Nerlov, Soheil Meshinchi, Lucia Carbone, Brian J. Druker, Julia E. Maxson

**Affiliations:** Knight Cancer Institute, Oregon Health & Science University, Portland, Oregon, 97239, USA; Division of Hematology and Medical Oncology, Oregon Health & Science University, Portland, Oregon, 97239, USA; Knight Cardiovascular Institute, Oregon Health & Science University, Portland, Oregon, 97239, USA; Division of Bioinformatics and Computational Biology, Oregon Health & Science University, Portland, Oregon, 97239, USA; Program in Molecular and Cellular Biology, Oregon Health & Science University, Portland, Oregon, 97239, USA; Computational Biology Program, Oregon Health & Science University, Portland, Oregon, 97239, USA; Department of Molecular and Medical Genetics, Oregon Health & Science University, Portland, Oregon, 97239, USA; Howard Hughes Medical Institute, Portland, OR, USA; Clinical Research Division, Fred Hutchinson Cancer Research Center, Seattle WA.; Division of Pediatric Hematology/Oncology, University of Washington, Seattle WA.; MRC Molecular Haematology Unit, MRC Weatherall Institute of Molecular Medicine, University of Oxford, Oxford, UK.

**Keywords:** Acute Myeloid Leukemia, Enhancers, CEBPA, CSF3R, Chromatin

## Abstract

Acute Myeloid Leukemia (AML) develops due to the acquisition of mutations from multiple functional classes. Here, we demonstrate that activating mutations in the granulocyte colony stimulating factor receptor (CSF3R), cooperate with loss of function mutations in the transcription factor CEBPA to promote acute leukemia development. This finding of mutation-synergy is broadly applicable other mutations that activate the JAK/STAT pathway or disrupt CEBPA function (i.e. activating mutations in JAK3 and Core Binding Factor translocations). The interaction between these distinct classes of mutations occurs at the level of myeloid lineage enhancers where mutant CEBPA prevents activation of subset of differentiation associated enhancers. To confirm this enhancer-dependent mechanism, we demonstrate that CEBPA mutations must occur as the initial event in AML initiation, confirming predictions from clinical sequencing data. This improved mechanistic understanding will facilitate therapeutic development targeting the intersection of oncogene cooperativity.

## INTRODUCTION

Acute Myeloid Leukemia (AML) is a deadly hematologic malignancy that results from the stepwise acquisition of genomic aberrations. The mutations that collaborate to produce AML often occur in distinct functional categories^1^. Class I mutations activate signaling pathways and in isolation produce disease with a myeloproliferative phenotype. Class II mutations perturb the function of transcription factors or epigenetic modifiers and alone can cause a myelodysplastic phenotype. However, when present in combination, Class I and Class II mutations produce a highly proliferative disease with a block in differentiation, both hallmarks of AML.

The transcription factor CCAAT enhancer binding protein alpha (CEBPA) is the master regulator of myeloid lineage commitment. CEBPA is recurrently mutated in AML and is a classic example of a class II mutation^2^. Mutations in CEBPA cluster at the N- and C-terminus of the protein. N-terminal mutations disrupt CEBPA-dependent cell cycle regulation while C-terminal mutations occur in the DNA-binding domain and produce dominant negative loss of function. Recent studies have identified a high rate of co-occurrence of mutations in CEBPA with mutations the granulocyte-colony stimulating factor receptor (CSF3R)^3–6^. CSF3R mutations most commonly occur in the membrane proximal region, and lead to ligand independent receptor dimerization and constitutive signaling via the JAK/STAT pathway. Similar to other Class I mutations, membrane proximal CSF3R mutations produce a myeloproliferative phenotype when present in isolation and are the major oncogenic driver of the disease chronic neutrophilic leukemia^7^. During normal myeloid development, G-CSF signaling via CSF3R leads to proliferation of myeloid precursors and neutrophilic differentiation. CEBPA is required for CSF3R-mediated transcription of myeloid specific genes, and myeloid differentiation arrests at the level of the common myeloid progenitor when CEBPA is deleted^8,9^. In patients harboring both CEBPA and CSF3R mutations, prognosis is worse than when mutant CEBPA is present in isolation^10^. In spite of the established functional interdependence of CSF3R and CEBPA during normal hematopoiesis, the mechanism by which oncogenic mutations in these two genes drive AML remains unknown.

Herein, we demonstrate that CSF3R and CEBPA cooperate to produce a highly proliferative immature myeloid leukemia in mice that phenocopies the human disease transcriptionally and morphologically. We establish that mutant CSF3R exerts much of its activity through the binding of the STAT family of transcription factors to promoters. In contrast mutant CEBPA exerts little oncogenic activity at promoters, instead impacting the activation of differentiation associated enhancers. We further demonstrate that mutant CEBPA must occur prior to mutant CSF3R in order for AML to initiate. This confirms predictions based on clinical sequencing and provides a novel model to study oncogene order.

## RESULTS

### CSF3R^T618I^ and CEBPA Mutations Cooperate to Produce Rapid Cell Growth

We decided to study a representative N-terminal and C-terminal mutation (F82fs and V314VW), in combination with CSF3R mutations, which frequently co-occur in AML^4^. When expressed in mouse bone marrow cells via retroviral transduction, neither CEBPA^F82fs^ nor CEBPA^V314VW^ produced colonies in cytokine free methylcellulose (Figure 1A, B). As previously reported, CSF3R^T618I^ produced a modest number of colonies in isolation^7^. The addition of CEBPA^V314VW^, but not CEBPA^F82fs^, dramatically augmented CSF3R^T618I^-driven colony production, produced indefinite replating, and supported sustained growth in liquid media lacking cytokines (Figure 1C). These results were confirmed with a second C-terminal CEBPA mutation CEBPA^K313KR^ (Figure 1D). To determine how co-expression of both N- and C-terminal CEBPA mutations would alter the effects of CSF3R^T618I^, we expressed all three mutations simultaneously. In this context, the addition of CEBPA^F82fs^ mildly augmented CFU formation beyond that seen with CSF3R^T618I^ and CEBPA^V314VW^ (Figure 1E, F). To determine whether this phenomenon was generalizable, we looked for other mutations that co-occur with CEBPA in AML. We identified a patient from a recently published data set with CEBPA-mutant AML who also had a previously characterized activating Class I mutation in JAK3 (JAK3^M511I^)^11,12^. Simultaneous introduction of these mutations promoted cytokine independent colony growth *in vitro* (Figure 1G,H). CSF3R mutations also co-occur with the chromosomal translocation t(8;21), leading to expression of the AML-ETO oncoprotein^4^. Similar to mutations in CEBPA, AML-ETO also augmented CSF3R^T618I^ - induced colony formation (Figure 1I, J). AML-ETO is known to down-regulate CEBPA, possibly providing a unifying mechanism for oncogenic cooperation with activated JAK/STAT signaling^13^. Consistent with this, we found that expression of AML-ETO in mouse bone marrow resulted in a decrease in CEBPA expression (Figure 1 K). Furthermore, bone marrow from CEBPA knockout mice transduced with CSF3R^T618I^ produced significantly more colonies than littermate control bone marrow (Figure L, M).

**Figure 1.**
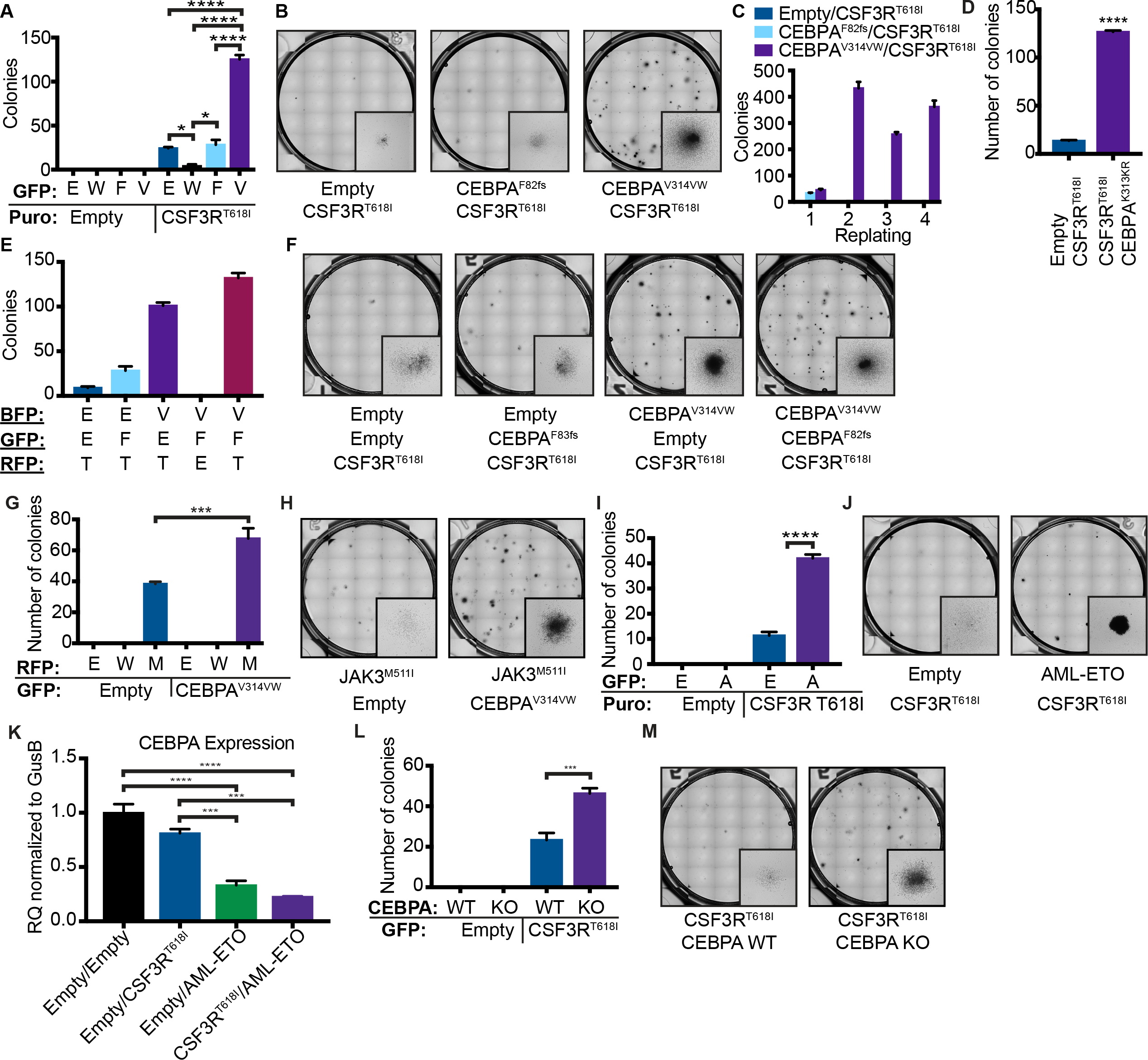
CSF3R^T618I^ and CEBPA Mutations Cooperate to Produce a Rapidly Lethal Leukemia. **A.** Colony assay from mouse bone marrow transduced with CSF3R^T618I^ and either CEBPA^WT^ (W), CEBPA^F82fs^ (F) or CEBPA^V314VW^ (V) (n=3/group). **B.** Representative images for A. **C.** Serial replating of colony assay in A. **D.** Colony assay from mouse bone marrow transduced with CSF3R^T618I^ and either empty vector or CEBPA^K313KR^. **E.** Colony forming assay from mouse bone marrow transduced with the combination of CSF3R^T618I^ (T), CEBPA^F82fs^ (F), CEBPA^V314VW^ (V), two oncogene combinations and empty vector controls. 1700 cells were plated per condition. **F.** Representative images for D. **G**. Colony assay from mouse bone marrow transduced with Empty Vector (E) JAK3^WT^ (W), JAK3^M511I^ (M), or CEBPA^V314VW^. **H.** Representative images of colony assay for G. **I.** Colony assay from mouse bone marrow transduced with Empty Vector (E), AML-ETO or CSF3R^T618I^. **J.** Representative colony assay images for I. **K.** CEBPA expression in mouse bone marrow transduced with Empty Vector (E), AML-ETO or CSF3R^T618I^ as measured by real time PCR and normalized to *GusB*. **L.** Colony assay from mouse bone marrow from CEBPA KO or Cre-littermate control bone marrow transduced with Empty Vector or CSF3R^T618I^. **M.** Representative colony assay images for L. In all cases n=/3 group and values are represented as mean +/− SEM (*: p<0.05, **: p<0.001, ***: p=0.0001, ****: p<0.001). Significance of comparisons assessed by ANOVA with Sidak post-test.

To characterize gene expression changes driven by CSF3R^T618I^ and CEBPA^V314VW^, we performed RNA-seq on mouse bone marrow transduced with CSF3R^T618I^, CEBPA^V314VW^, or the combination of both. We used a linear model with an interaction term (Figure 2A), and identified 683 genes that were significantly up or down regulated in response to CSF3R^T618I^ (q <0.05, Log_2_ fold change >1 or <−1). We also identified 809 genes up or down regulated in response to CEBPA^V314VW^ expression. Additionally, there were 570 genes that demonstrated an interaction between CSF3R^T618I^ and CEBPA^V314VW^ (effect less or more than additive). To identify enriched transcriptional programs, we performed Gene Set Enrichment Analysis (GSEA) comparing CSF3R^T618I^ to all other conditions (Figures 2B, Figure S1A). CSF3R^T618I^ dramatically up-regulated genes associated with myeloid differentiation, while CEBPA^V314VW^ strongly down-regulated them. In addition, genes associated with the wild type CEBPA network followed a similar pattern (Figure 2C). This suggests that myeloid differentiation in response to CSF3R^T618I^ is, at least in part, dependent on CEBPA.

**Figure 2.**
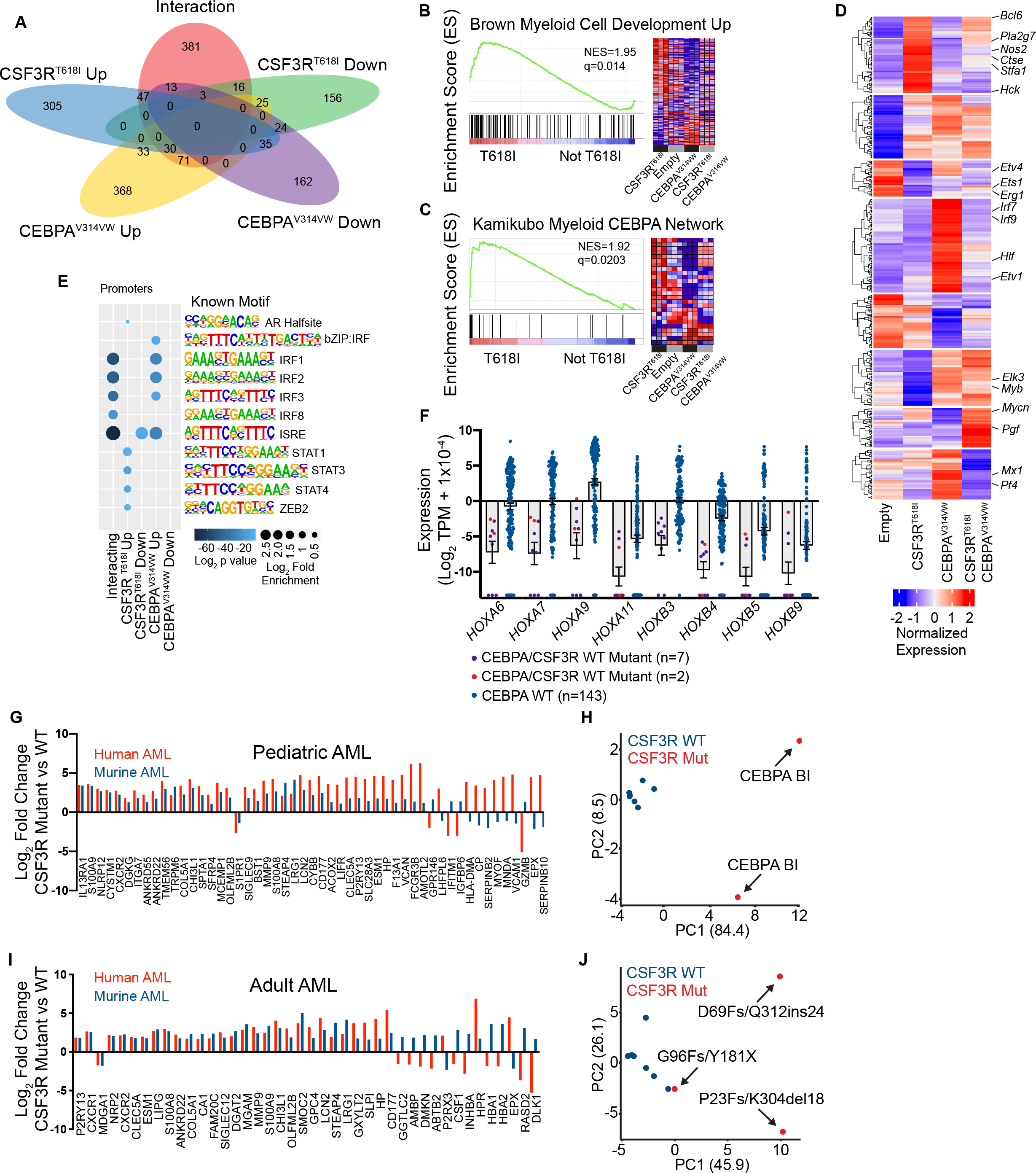
Mutant CEBPA Blocks Myeloid Differentiation in Response to CSF3R. **A.** Venn diagram of differentially expressed genes from RNA-seq on lineage negative mouse bone marrow transduced with empty vector, CSF3R^T618I^, CEBPA^V314VW^ or the combination of oncogenes. **B, C.** Select enriched gene sets in CSF3R^T618I^ vs. other categories NES= normalized enrichment score (q<0.05). **D.** Hierarchical clustering of interacting genes. **E.** Motif enrichment at promoters of differentially expressed (DE) genes. Top 5 motifs per category with q<0.05 shown. **F.** Expression of differentially expressed HOX genes in pediatric AML patients harboring CEBPA mutations. **G.** Expression of genes differentially expressed in both murine and pediatric human CSF3R/CEBPA AML as compared with CEBPA-mutant CSF3R-WT AML. **H.** PCA analysis of pediatric CEBPA mutant AML using convergent human-mouse gene set. **I**. Expression of genes differentially expressed in both murine and adult human CSF3R/CEBPA AML as compared with CEBPA-mutant CSF3R-WT AML. **J.** PCA analysis of adult CEBPA mutant AML using convergent human-mouse gene set.

To further describe the transcriptional changes associated with concurrent oncogene expression, we performed hierarchical clustering of the interacting genes (Figure 2D & Table S1). Of particular interest was a cluster of genes that were strongly up-regulated by CSF3R^T618I^, but suppressed by co-expression of CEBPA^V314VW^, as this mirrored the pattern of myeloid differentiation. This cluster contained genes such as *Nos2*, *Hck*, *Stf1a*, and *Pla2g7*, all of which are expressed in mature neutrophils^14^. To determine, whether CEBPA binding was significantly associated with these interacting genes, we analyzed published ChIP-seq data from mouse granulocyte-macrophage progenitors (GMPs) ^15^ and showed that 60% of the interacting genes were within 2 Kb of a CEBPA peak, compared with approximately 20% of non-interacting genes (Figure S1C). We next performed motif enrichment analysis at the promoters of each category of differentially expressed genes (Figures 2E & S1B). This identified a strong enrichment of STAT binding sites in the promoters of CSF3R^T618I^ up-regulated genes. Surprisingly however, we did not detect CEBPA motif enrichment at the promoters of any of the groups of DE genes, suggesting that CEBPA^V314VW^ may impact CSF3R^T618I^-induced myeloid differentiation through binding to non-promoter regulatory regions.

To validate these findings in human AML, we examined RNA-seq data from the pediatric TARGET initiative^16^. A total of 152 patients were evaluated, including seven patients with CEBPA mutations, and 2 patients with both CSF3R and CEBPA mutations. Differential gene expression revealed markedly decreased HOX gene expression in CEBPA mutant samples compared with CEBPA WT samples, a well-established finding in adult CEBPA mutant AML (Figure 2F, Table S2)^17^. Interestingly, CSF3R mutant samples tended to display increased HOX gene expression although this did not rise to the level of statistical significance. Comparison of CEBPA mutant/CSF3R WT and CEBPA mutant/CSF3R mutant patient samples revealed numerous differentially-expressed genes (Table S2). Principle component analysis using these differentially-expressed genes clearly separates CEBPA/CSF3R mutant samples from those harboring mutant CEBPA alone (Figure S1D). Comparison of these differentially-expressed genes to those identified in mouse revealed 52 ortholog pairs with differential expression driven by mutant CSF3R (Figure 2G). Of these, 75% demonstrated concordant regulation between mouse and human. A similar analysis was performed on adult CEBPA/CSF3R mutant AML samples from the Leucegene cohort, which also demonstrated that the preponderance of orthologous gene pairs display concordant regulation (Figure 2I) ^3^. Both human/mouse concordant gene sets were able to independently segregate CSF3R mutant from wild type samples by PCA (Figure 2H, J).

### JAK/STAT Activation and CEBPA Dysregulation Cooperate to Promote Myeloid Leukemia *in vivo*

To establish whether oncogenic CSF3R and CEBPA mutations collaborate *in vivo* to produce AML, we performed murine bone marrow transplantation with retrovirally introduced CSF3R^T618I^ alone or in combination with CEBPA^V314VW^ or CEBPA^F82fs^. Mice receiving cells transduced with CSF3R^T618I^ and CEBPA^V314VW^ developed a myeloid leukemia that was uniformly lethal by day 14 post-transplant and was associated with leukocytosis and marked splenomegaly (Figures 3A-B, F-G & Figure S2A). In the bone marrow, normal hematopoiesis was completely replaced by large cells with myeloblastic morphology (Figures 3C, S2B-D). The blasts were positive for CD11b, but were dim for GR-1 relative to the CD11b+/GR−1^high^ cells seen in CSF3R^T618I^ alone mice sacrificed at an early time point (Figure 3D). The CSF3R/CEBPA mutant leukemia was also amenable to serial transplant in up to quaternary recipients (Figure S3A-C). Mice harboring CSF3R^T618I^ alone or in combination with CEBPA^F82fs^ developed disease with a long latency, associated with leukocytosis, variable splenomegaly, mature immunophenotype, and morphologic neutrophils in the peripheral blood and bone marrow (Figures 2A-B, & S3D-H). Similarly, the combination of JAK3^M511I^ with CEBPA^V314VW^ also produced a rapidly lethal myeloid leukemia with markedly reduced latency compared with expression of either mutation alone (Figure 3H-J). In addition, the combination of AML-ETO and CSF3R^T618I^ also produced an accelerated undifferentiated myeloid leukemia (Figure 3K-M). Collectively, these data show that activation of JAK-STAT signaling and disruption of CEBPA are sufficient to drive leukemia development with all the cardinal features of AML.

**Figure 3.**
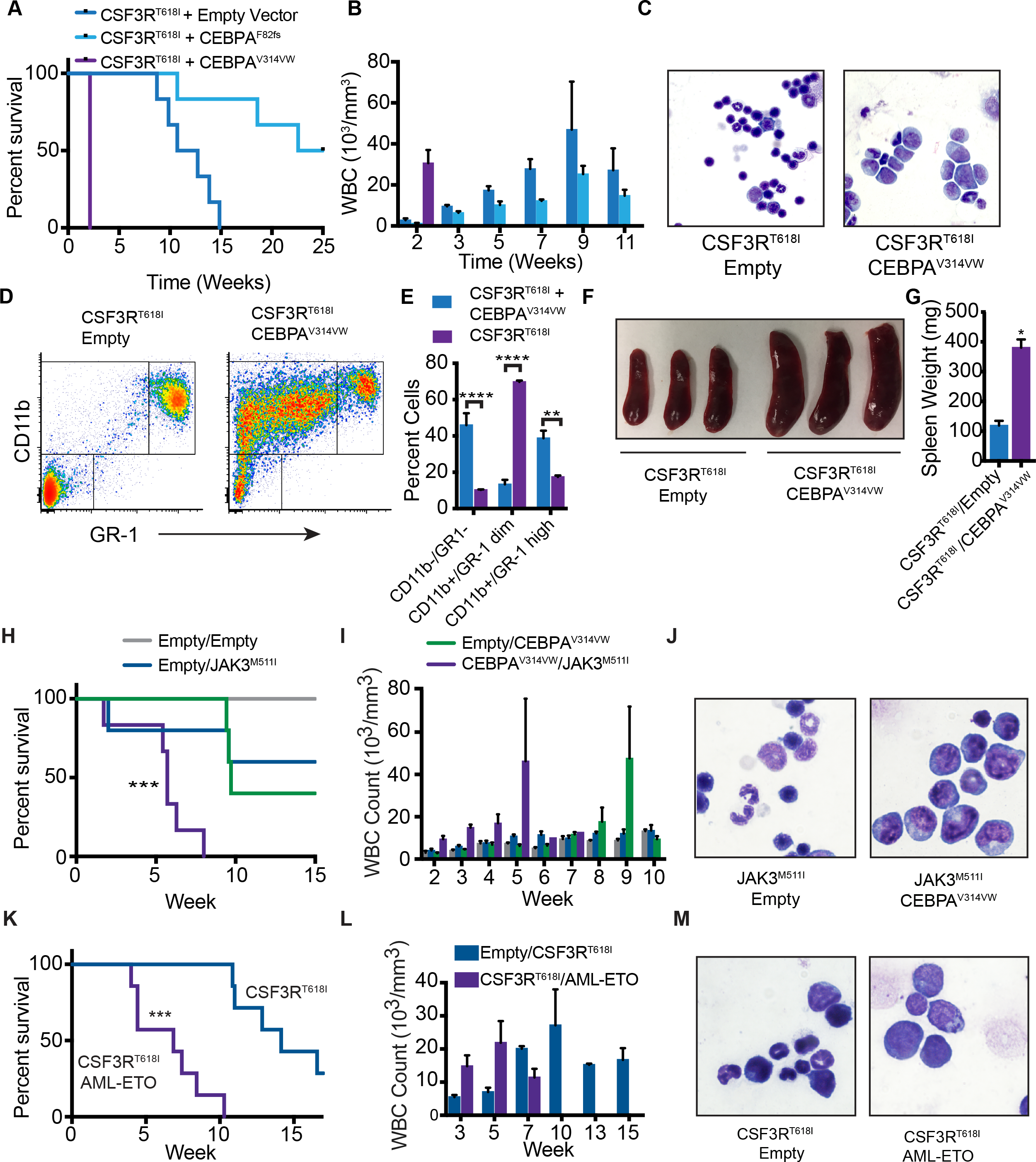
JAK/STAT Activation and CEBPA Dysregulation Cooperate to Promote Myeloid Leukemia *in vivo.* **A.** Survival of mice transplanted with 100,000 cells containing CSF3R^T618I^ + Empty vector, CSF3R^T618I^ + CEBPA^F82fs^, or CSF3R^T618I^ + CEBPA^V314VW^ (n=6/group). All three pairwise comparisons were significant by ANOVA. **B.** WBC counts of mice harboring mutation combinations in **A** (n= 6/group at experiment start)**. C.** Bone marrow smears (representative of 3 animals). **D.** Bone marrow flow cytometry for GR-1 and CD11b from mice transplanted with 10,000 cells containing CSF3R^T618I^ + Empty vector, or CSF3R^T618I^ + CEBPA^V314VW^ sacrificed on day 18 post-transplant. **E.** Quantification of cell populations from **D** (n=3/group). **F.** Representative images of spleens from groups in **C**. **G.** Spleen weights from groups in **C** (n=3/group). **H.** Survival of mice transplanted with 100,000 cells containing Empty Vector, JAK3^M511I^ and CEBPA^V314VW^ with empty vector or both oncogenes in combination (n=6/group). **I.** WBC counts. **J.** Representative bone marrow histology from mice in **H** at time of survival endpoint (representative of 3 animals/group). **K.** Survival of mice transplanted with 10,000 cells containing CSF3R^T618I^ + empty vector or CSF3R^T618I^ +AML-ETO (n=7/group). **L.** WBC counts. **M.** Bone marrow smears (representative of 3 mice per group). In all cases, values are represented as mean +/− SEM (*: p<0.05, **: p<0.001, ***: p=0.0001, ****: p<0.001.) Survival assessed by Log Rank test. Significance of other comparisons assessed by Student’s t-test for two group comparisons or ANOVA with Sidak post-test, as appropriate.

### CEBPA Mutations Disrupt Activation of Myeloid Lineage Enhancers

During normal hematopoietic development, CEBPA is responsible for establishing the enhancer landscape that permits myeloid differentiation^18^. We therefore hypothesized that CEBPA^V314VW^ blocks differentiation by inhibiting priming or activation of myeloid lineage enhancers. To test this, we utilized murine HoxB8-ER cells, which mimic GMPs and differentiate down the neutrophilic lineage to become CD11b and GR-1 positive after estrogen withdrawal^19^. In this model, expression of CSF3R^T618I^ accelerated differentiation, while CEBPA^V314VW^ blocked differentiation, as measured by CD11b and GR-1 expression (Figures 4A & S4A). The combination of mutations produced maturation arrest, similar to CSF3R/CEBPA mutant murine leukemia. We confirmed that neither oncogene was changing the expression of the other by qPCR (Figure S4B). A similar differentiation-block was seen when AML-ETO was co-expressed with CSF3R^T618I^ (Figure S4C). Thus, HoxB8-ER cells expressing CSF3R^T618I^ and CEBPA^V314VW^ recapitulate the phenotype seen in our *in vivo* model.

**Figure 4.**
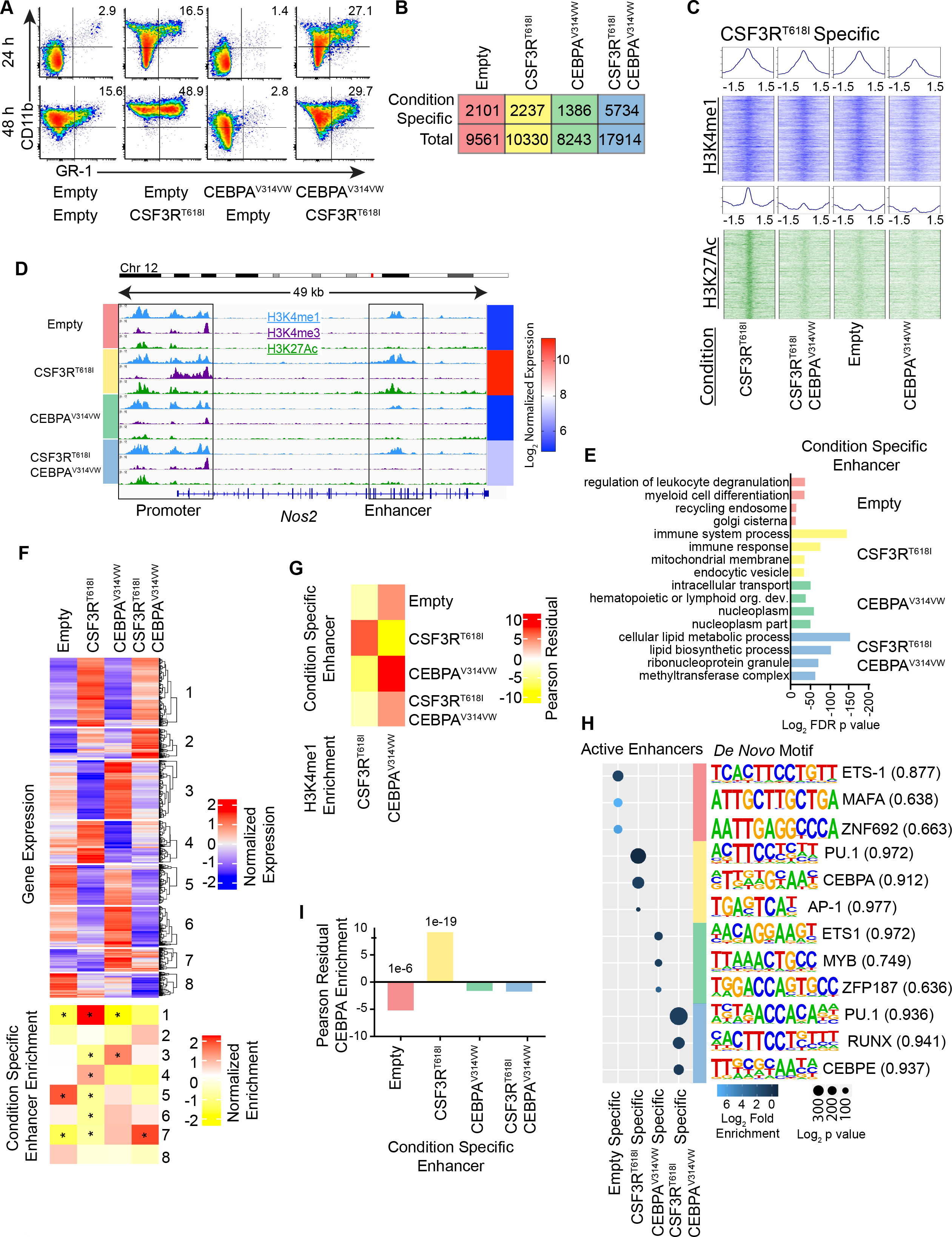
CEBPA Mutations Disrupt Activation of Myeloid Lineage Enhancers. **A.** Expression of CD11b and GR-1 by flow cytometry in murine HoxB8-ER cells transduced with empty vector, CSF3R^T618I^, CEBPA^V314VW^ or the oncogene combination 24-48 hours after estrogen withdrawal. **B.** Number of condition specific enhancers identified in this study. **C**. ChIP-seq heat maps for H3K4me1 and H3K27ac at +1.5 Kb around the center of CSF3R^T618I^-specific enhancers, across treatment conditions. ChIP tracks display fold enrichment relative to corresponding input. **D.** The epigenetic landscape at the *Nos2* locus demonstrates condition-specific promoter and enhancer activation. **E.** Gene ontology analysis for condition specific enhancers. **F.** Unsupervised hierarchical clustering of differentially expressed genes (as assessed by microarray) and associated enrichment analysis for condition specific enhancers. Bottom heat map displays normalized Pearson residual values from χ^2^ test for each category of condition specific enhancer and gene expression cluster. Cluster numbers appear on the right side and *: p<0.05 adjusted for multiple comparisons. **G.** Association of H3K4me1 enriched peaks with condition specific enhancers. Heat map displays normalized Pearson residual values from χ^2^ test for each category of condition specific enhancer and gene expression cluster. All comparisons have p<0.05 adjusted for multiple comparisons made by the method of Holm-Bonferroni. **H.** *De novo* motif enrichment in condition specific enhancers. Values in parenthesis correspond to motif match score. **I**. Assessment of CEBPA ChIP-seq peak overlap with condition specific enhancers by χ^2^ test with Pearson residual values plotted. Adjusted p values (by the method of Holm-Bonferroni) are displayed above bars.

To study enhancer dynamics in the context of these mutations, we performed chromatin immunoprecipitation sequencing (ChIP-seq) for H3K4me1, H3K4me3 and H3K27ac in HoxB8-ER cells transduced with CSF3R^T618I^, CEBPA^V314VW^, both mutations, or empty vectors. We identified 42,124 active enhancers (defined as high H3K4me1, low H3K4me3 and high H3K27ac) across all 4 conditions (Table S4). To validate these findings in human AML, we mapped our enhancers to orthologous human genomic coordinates and assessed overlap of enhancers only active in the CSF3R^T618I^/CEBPA^V314VW^ condition with those present in human AML with mutant CEBPA^20^. We observed a significant enrichment of overlap between our mouse AML enhancers with those found in human AML, suggesting conserved biology (Figure S5D). We performed a similar analysis using regions of DNase I hypersensitivity from human CEBPA mutant AML and again saw marked enrichment (Figure S5E).

We next focused on enhancers that were active exclusively in one condition (Condition-Specific Enhancers, Figures 4B, C & S5A-C). An example of a CSF3R^T618I^-specific enhancer in proximity to the *Nos2* gene is shown (Figure 4D). Gene ontology analysis on the 2,237 CSF3R^T618I^-specific enhancers revealed enrichment for terms associated with immune responses and phagocytosis, demonstrating that these genes are associated with the mature neutrophil phenotype (Figure 4E, Table S5). To understand whether CEBPA^V314VW^ blocks myeloid differentiation through disruption of CSF3R^T618I^-specific enhancers and associated gene expression, we performed microarray gene expression analysis on HoxB8-ER cells harboring each oncogene in isolation or the combination (Figure 4F, Table S3). The expression profiles of key genes were validated by qPCR (Figure S5B). Genes associated with mature myeloid phenotypes, such as *Nos2*, *Hck, Bcl6* and *Pla2g7*, displayed increased expression in the CSF3R^T618I^ condition, and were repressed with co-expression of CEBPA^V314VW^. To globally assess whether CSF3R target genes were associated with CSF3R^T618I^-specific enhancers, we performed hierarchical clustering on genes that were differentially expressed in one or more treatment condition. We then examined enrichment of each condition-specific enhancer group across these clusters (Figure 4F, Table S3). Specifically, we observed strong enrichment of CSF3R^T618I^-specific enhancers in Clusters 1 and 4, which displayed the highest level of gene expression with CSF3R^T618I^ alone.

In contrast with enhancer activation, there was little difference in enhancer priming (as denoted by presence/absence of H3K4me1 peaks) across the four treatment conditions. However, the intensity of H3K4me1 signal at CSF3R^T618I^-specific enhancers varied between conditions. H3K4me1 enrichment at CSF3R^T618I^-specific enhancers was strongest in cells expressing CSF3R^T618I^ only, and lowest in cells expressing CEBPA^V314VW^ only (Figure 4G). To quantify this, we identified regions with significantly different H3K4me1 enrichment between CSF3R^T618I^ and CEBPA^V314VW^ conditions. This analysis revealed 798 regions of H3K4me1 that were specific to CSF3R^T618I^ and 833 regions specific to CEBPA^V314VW^. Regions with higher H3K4me1 enrichment in CSF3R^T618I^ were highly associated with enhancers that were specifically activated by CSF3R^T618I^. In contrast, regions with higher H3K4me1 enrichment in the CEBPA^V314VW^ condition were associated with CEBPA^V314VW^-specific enhancers. This demonstrates that in addition to regulating enhancer activation, CSF3R^T618I^ and CEBPA^V314VW^ also affect H3K4me1 levels.

To identify possible transcriptional regulators, we performed transcription factor motif enrichment. This analysis demonstrated enrichment of CEBPA motifs in CSF3R^T618I^-specific enhancers, but not in any other group (Figure 4H). This suggests that CEBPA^V314VW^ blocks differentiation by disrupting the interaction of wild type CEBPA with CSF3R^T618I^-specific enhancers. To provide further evidence for this hypothesis, we utilized published CEBPA ChIP-seq data from GMPs (the nearest normal cell to HoxB8 immortalized progenitors)^15^. We found strong enrichment of CEBPA peaks overlapping CSF3R^T618I^-specific enhancers, but not the other condition-specific enhancer groups (Figure 4I). These data support a model in which CEBPA^V314VW^ displaces wild type CEBPA from CSF3R^T618I^-specific enhancers, preventing enhancer activation and subsequent expression of differentiation-associated genes.

### CEBPA Mutations Must Precede Mutations in CSF3R for AML Initiation

Enhancer priming and activation precede promoter activation, suggesting that epigenetic changes at enhancers must occur as early events^21^. Clinical sequencing data demonstrates that CEBPA mutations frequently occur at higher variant allele frequencies than CSF3R mutations^3,4^. This has led to the prediction that CEBPA mutations occur early in disease development. Conversely however, it is possible that CEBPA mutations could be acquired late during blast-crisis transformation of chronic neutrophilic leukemia. To evaluate the impact of mutation order on AML development, we developed a Cre-inducible retroviral vector. We paired this vector with hematopoietic cells from Rosa 26^ERT2^-Cre, which is activated via tamoxifen administration (Figure 5A). We co-expressed this inducible vector with a constitutive vector, enabling the study of two distinct mutational orders of acquisition: CSF3R^T618I^ first and CEBPA^V314VW^ first. We first assessed the impact of mutation order on bone marrow colony formation by plating both ordered combinations in the presence of 4-hydroxytamoxifen (4-OHT), thus delaying the expression of the second oncogene by approximately 48 hours. Strikingly, we found that CSF3R^T618I^-first produced far fewer colonies than CEBPA^V314VW^-first (Figure 5B, C). Numerically and morphologically, CSF3R^T618I^-first colonies were closer to CSF3R^T618I^-only. This finding was not due to differential expression of CEBPA from the inducible and constitutive vectors, as both showed equivalent CEBPA expression (Figure 5D).

**Figure 5.**
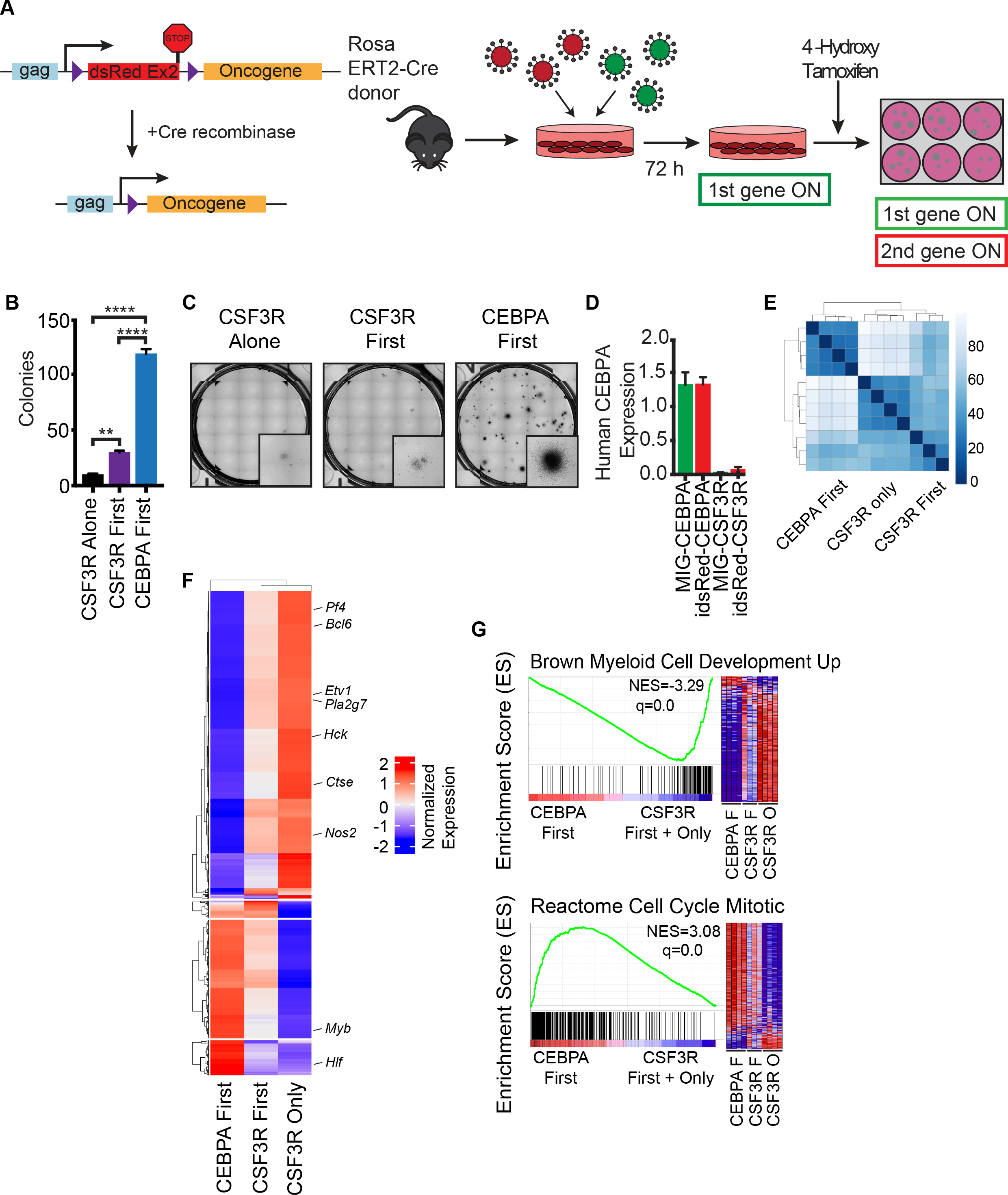
CEBPA Mutations Must Precede Mutations in CSF3R to Block Differentiation. **A**. Diagram of order of acquisition system. Oncogenes in the MIG (GFP+) vector tagged with GFP are constitutively expressed, and oncogenes in the idsRed vector are expressed only after Cre mediated recombination. Expression of the second oncogene can be induced by culture in 4-OHT. **B.** Colony assay from mouse bone marrow transduced with CEBPA First (MIG-CEBPA^V314VW^ +idsRed-CSF3R^T618I^) CSF3R First (MIG-CSF3R^T618I^ + idsRed-CEBPA^V314VW^) and CSF3R Only (MIG-CSF3R^T618I^ + idsRed Empty) and plated in 4-OHT (n=3/group). **: p<0.01, ****: p<0.0001, as measured by ANOVA with Sidak’s post-test correction. **C.** Human CEBPA and CSF3R expression in mouse bone marrow expressing MIG-CEBPA^V314VW^, idsRed-CEBPA^V314VW^, MIG-CSF3R^T618I^ or idsRed-CSF3R^T618I^ measured by TaqMan quantitative PCR (n=3/group). **D.** Representative images of colony assay. **E.** Clustering by Euclidian distance for RNA sequencing performed on lineage negative mouse bone marrow expressing CEBPA-first, CSF3R-first or CSF3R-only (n=3-4/group). **F.** Unsupervised hierarchical clustering of top 750 differentially expressed genes per pairwise comparison (q<0.05, log_2_Fold Change <−1 or >1) (n=3-4/group). **G.** Representative enriched gene sets from GSEA performed on samples from **E** (n=3-4/group). In all cases, values are represented as mean +/− SEM.

To establish the transcriptional profile of these two distinct mutation orders, we performed RNA-seq on mouse bone marrow transduced with CSF3R^T618I^-only, CSF3R^T618I^-first, and CEBPA^V314VW^-first, cultured for 48 hours with 4-OHT. Unsupervised clustering by Euclidean distance revealed that CSF3R^T618I^-only and CSF3R^T618I^-first cluster together and are globally distinct from CEBPA^V314VW^-first (Figure 5E). The expression of differentiation-associated genes (*Nos2*, *Hck*, *Bcl6*, *Pla2g7*) in CSF3R^T618I^-first cells was intermediary between CSF3R^T618I^-only and CEBPA^V314VW^-first (Figure 5F, Table S6). GSEA revealed enrichment for numerous signatures associated with myeloid differentiation and inflammatory responses in CSF3R^T618I^-first/only cells (Figures 5G & S6). CEBPA^V314VW^-first cells demonstrated enrichment for signatures associated with cell cycle progression, Myc signaling, and stem/progenitor cell phenotype. When comparing the expression of gene sets across samples, CSF3R^T618I^-first cells were intermediary between CSF3R^T618I^-only and CEBPA^V314VW^-first. Taken together, these studies demonstrate a profound effect of mutation order on gene expression and hematopoietic phenotype *in vitro*.

We next explored the contribution of mutation order to disease latency and phenotype *in vivo*. Mice were transplanted with CSF3R^T618I^-first or CEBPA^V314VW^-first cells and allowed to recover for 4-weeks post-transplant. After 4-weeks, the second oncogene was induced with tamoxifen (Figure 6A). CEBPA^V314VW^-first mice succumbed to lethal myeloid leukemia with a median survival of 3.5 weeks post-tamoxifen. CEBPA^V314VW^-first leukemia was associated with splenomegaly and circulating blasts (Figure 6B, D, F, G, Figure S7 A-E). Bone marrow cells from CEBPA^V314VW^-first mice displayed a similar immature myeloid immunophenotype as mice transplanted with both oncogenes expressed simultaneously (Figure 6C). In contrast, CSF3R^T618I^-first mice displayed a differentiated myeloid immunophenotype when sacrificed at an early time point. Only one CSF3R^T618I^-first mouse developed leukemia at 7 weeks post-tamoxifen administration. Bone marrow cytology revealed a blast-like morphology, however flow cytometry revealed increased GR-1 staining, indicative of a higher degree of myeloid maturation (Figure 6C). Collectively, these data reveal that when CEBPA mutations are introduced after mutations in CSF3R, they are unable to fully block myeloid differentiation. Importantly, this impaired ability to block differentiation disrupts the development of AML *in vivo*.

**Figure 6.**
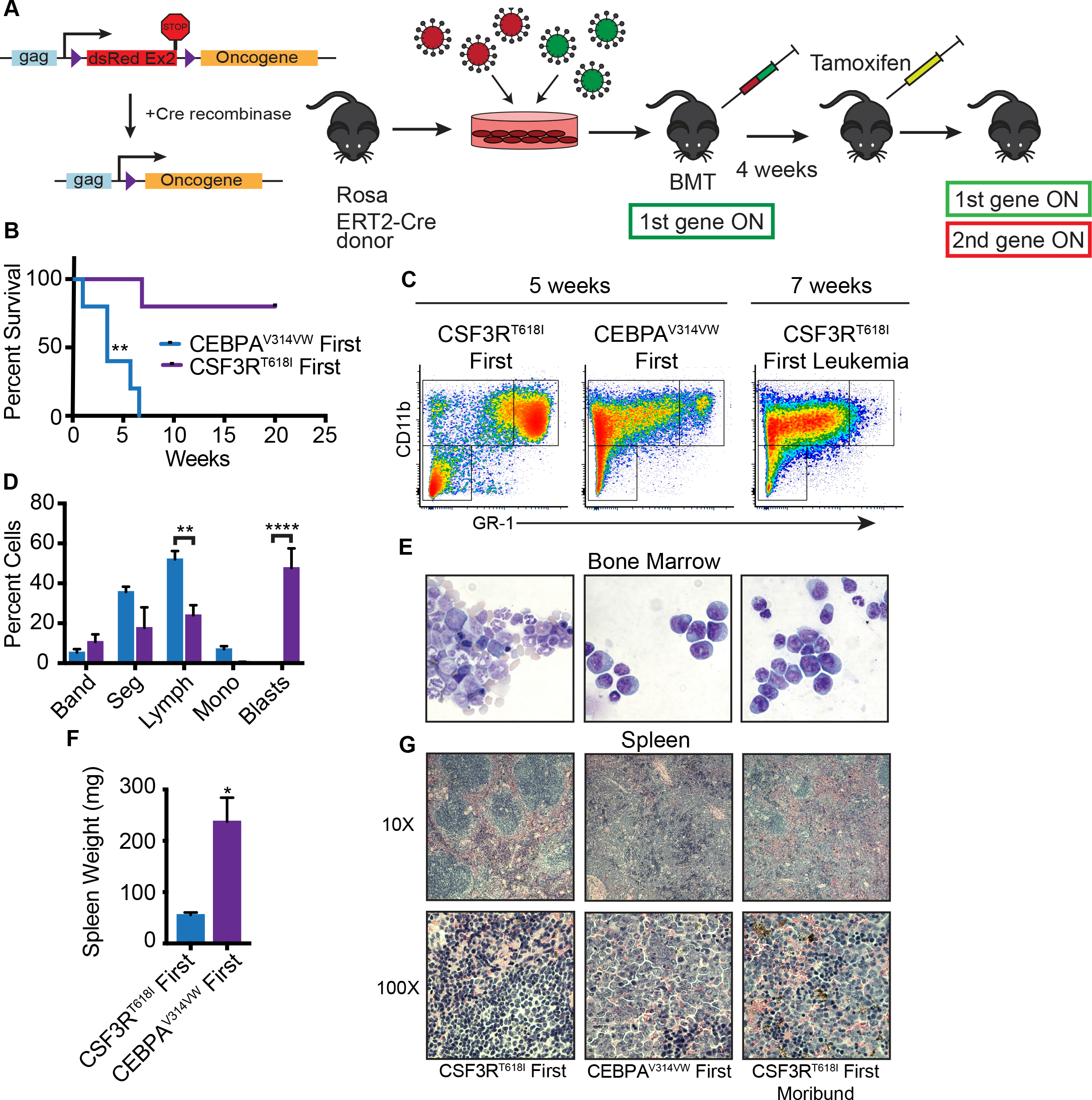
CEBPA Mutations Must Precede Mutations in CSF3R For AML Initiation. **A**. Diagram of order of acquisition system. Oncogenes in the MIG vector tagged with GFP are constitutively expressed and oncogenes in the idsRed vector are expressed only after Cre mediated recombination. Expression of the second oncogene can be induced by administration of tamoxifen. Mice were administered 75,000 double fluorescent cells. **B.** Survival reported as time from day one of tamoxifen induction (n=5/group). Statistical significance calculated by Log Rank test, **: p<0.01. **C.** Spleen weight of mice sacrificed at 5 weeks post tamoxifen treatment. **D.** Expression of CD11b and GR-1 in bone marrow at 5 weeks in moribund CEBPA^V314VW^ first mice, healthy CSF3R^T618I^ first mice or at 7-weeks in leukemic CSF3R^T618I^ first mice (only one animal represented by CSF3R first leukemia, others are representative of 3/group). **E.** Manual differentials of peripheral blood from mice 5 weeks after tamoxifen induction. **F.** Representative bone marrow histology. **G.** Representative spleen histology. In all cases, values are represented as mean +/− SEM.

## DISCUSSION

Our study adds to a growing body of data demonstrating that enhancer biology is integral to the development of hematologic malignancies. While AML is very heterogeneous from a genomic standpoint, recent work demonstrates that there are only a few epigenetic disease subtypes^20,26^. Thus, understanding the epigenetic dysfunction in AML may provide broad insight into therapeutic approaches. Although multiple global epigenetic regulators are recurrently mutated in AML, these have little impact on the organization of the epigenome^20,26^. Instead, mutations in lineage determining transcription factors are a major determinant of clustering. Using a DNase-I sequencing approach, Assi et al recently showed that core binding factor translocations and CEBPA mutant AML cluster together, consistent with our findings here of shared biology between AML-ETO and CEBPA^26^.

Although we focused on differentiation-associated enhancers driven by the CSF3R^T618I^, our study revealed a second set of enhancers that were activated exclusively in the presence of both mutant CSF3R and CEBPA. Both subsets of enhancers demonstrated strong enrichment of PU.1 motifs, consistent with the known role of this transcription factor in driving myeloid development. In normal hematopoiesis, PU.1 and CEBPA cooperate to open myeloid lineage enhancers with PU.1 performing pioneering function in early progenitors and CEBPA assuming this role late^18^. Thus, it is likely that certain early myeloid lineage enhancers can be activated by CSF3R^T618I^ in a CEBPA-independent manner via PU.1. Another interesting finding was the enrichment of RUNX motifs in enhancers activated only in the presence of both oncogenes. The RUNX family of transcription factors are critical to normal hematopoietic development, and in addition to being the targets of chromosomal translocations in core binding factor AML, also frequently harbor point mutations in AML^27^. Interestingly, Runx1 haploinsufficiency leads to hypersensitivity to GCSF, suggesting a negative feedback role^28^. The role of RUNX transcription factors in driving CEBPA/CSF3R mutant AML is an interesting area for future work.

We present the first direct evidence that the order in which oncogenic mutations occur is a major determinant of leukemia development (Figure 7G). Our finding that myeloid differentiation blockade can only occur with a distinct mutational order may also be a broadly conserved mechanism that applies to Class I and Class II mutation pairings. As nearly all prior studies have investigated mutation cooperation in the setting of simultaneous introduction, it is likely that important aspects of order-dependent disease biology have not yet been discovered. If reprograming of the lineage-specific enhancer repertoire is a common initiating event in AML, this creates even more impetus for the development of treatment strategies targeting these epigenetic pathways.

**Figure 7.**
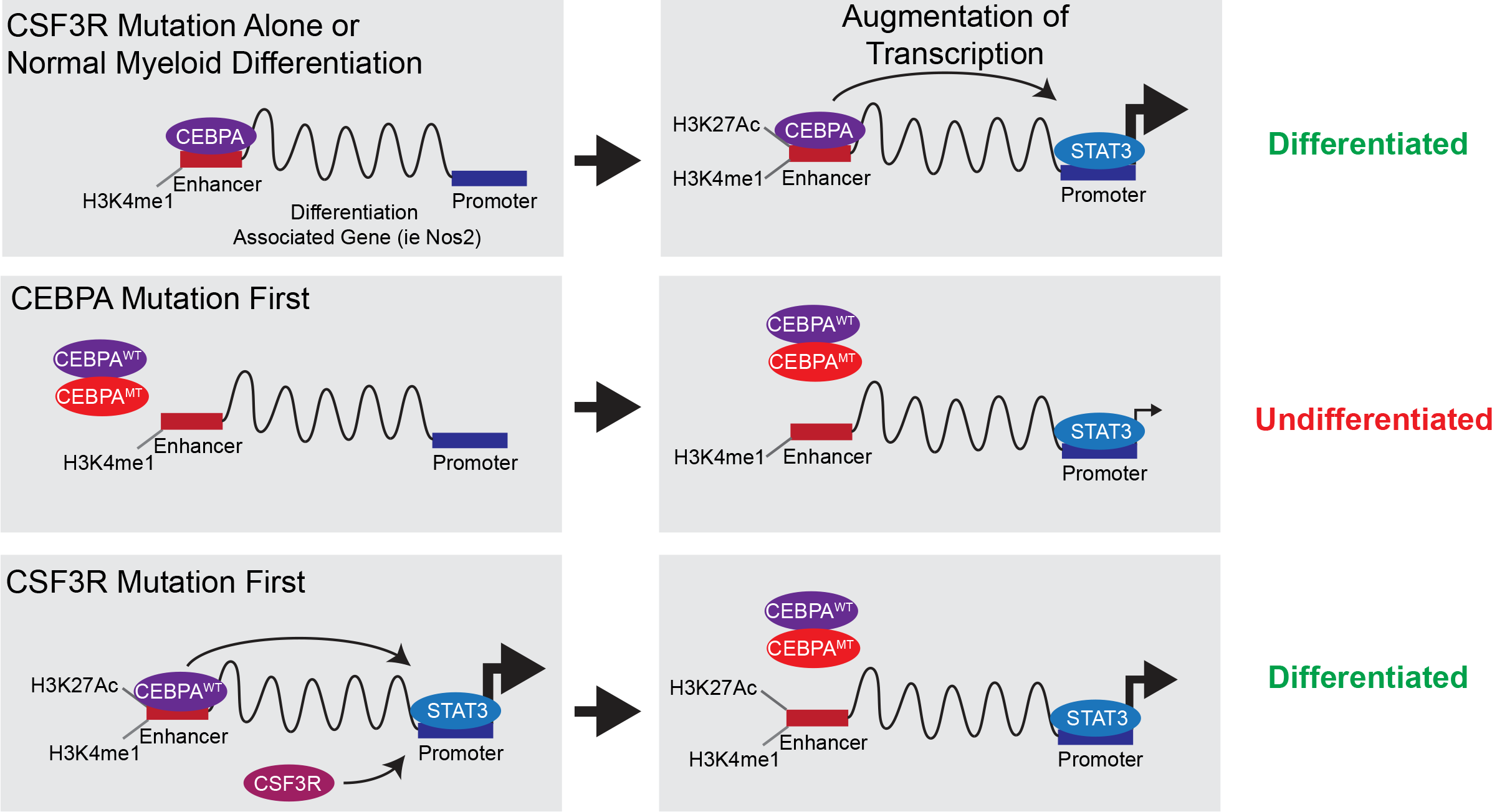
CEBPA Mutations Initiate AML Through Disruption of Myeloid Lineage Enhancers. During normal myeloid differentiation, CEBPA primes differentiation-associated enhancers leading to enhancer activation and subsequent augmentation of transcription of CSF3R/JAK/STAT target genes. Mutant CEBPA displaces WT CEBPA from these enhancers preventing activation of CSF3R target genes. When CSF3R mutations are introduced early, they activate transcription using the native enhancer repertoire initiating differentiation. Importantly this differentiation is insensitive to subsequent introduction of mutant CEBPA.

Our finding that epigenetic dysfunction is an obligate early event is likely to be generalizable to other forms of cancer. In renal cell carcinoma, loss of the tumor suppressor von Hippel-Lindau (VHL) is widely regarded as an initiating event and associated with dramatic changes in global DNA methylation, potentially impacting subsequently acquired signaling mutations ^22^. In breast cancer, mutation order is an important determinant of cancer phenotype, with luminal-type tumors demonstrating early loss of PTEN while basal-type tumors display early p53 mutation ^23^. In colon cancer, the canonical APC mutations decrease DNA methylation through upregulation of demethylases, which no doubt alter the impact of RAS mutations acquired later in disease evolution ^24^. Order-dependent mutational phenotypes were recently reported in myeloproliferative neoplasms, where the order of mutations in JAK2 and TET2 are important determinants of disease phenotype ^25^. When TET2 precedes JAK2, patients are more likely to present with polycythemia vera. In contrast, patients with JAK2-first disease were more likely to have essential thrombocytosis. Thus, it appears that preceding TET2 mutation provides an epigenetic context supporting erythroid development for subsequently-acquired JAK2 mutations.

In summary, we describe the epigenetic mechanism by which mutant CEBPA and CSF3R interact to drive AML development. Our study demonstrates that a subset of differentiation-associated enhancers are dysregulated by mutant CEBPA preventing normal myeloid maturation. Critically, this epigenetic dysregulation must occur as the initial event otherwise AML does not develop. These differentiation-associated enhancers represent a promising novel therapeutic target in this poor prognosis molecular subtype of AML.

## AUTHOR CONTRIBUTIONS

Conceptualization: T.P.B., B.J.D and J.E.M. Methodology: T.P.B., M.O. and J.E.M. Software: T.P.B, M.O, B.D. and S.J. and S.M. Formal Analysis: T.P.B, M.O., B.D. and S.J. Investigation: T.P.B, C.C., S.A.C, A.F., Z.S., K.N., D.L., B.G. and J.E.M. Data Curation: T.P.B, M.O, B.D. and S.J. and S.M. Writing – Original Draft: T.P.B., L.C. and J.E.M. Writing – Review & Editing: T.P.B, M.O., L.C., B.J.D. and J.E.M. Supervision: T.P.B., S.M., L.C., B.J.D. and J.E.M. Funding Acquisition: T.P.B., B.J.D., and J.E.M.

## ACKNOWLEDGEMENTS

Funding provided by an American Society of Hematology Research Training Award for Fellows and Collins Medical Trust Award to T.P.B., NCRR P51 OD011092 and KCVI to L.C. OHSU School of Medicine Faculty Innovation Fund to B.J.D. and J.E.M, Howard Hughes Medical Institute to B.J.D, NCI R00-CA190605, an ASH Scholar Award, and a MRF New Investigator Grant to J.E.M. We are grateful to the following core facilities and shared resources for their assistance with this work: OHSU Histopathology Shared Resource, OHSU Massively Parallel Sequencing Shared Resource, OHSU Gene Profiling Shared Resource, OHSU/KCVI Epigenetics Consortium, Knight Cancer Institute Biostatistics Shared Resource and the OHSU Flow Cytometry Shared Resource. We also thank the OHSU ExaCloud Cluster Computational Resource and the Advanced Computing Center that allowed us to perform the intensive large-scale data workflows.

We are grateful to the following core facilities: Histopathology Shared Resource, Massively Parallel Sequencing Shared Resource, Gene Profiling Shared Resource, Epigenetics Consortium, Knight Cancer Institute Biostatistics Shared Resource, Flow Cytometry Shared Resource, ExaCloud Cluster Computational Resource and the Advanced Computing Center.

## SUPPLEMENTAL FIGURE LEGENDS

**Figure S1.**
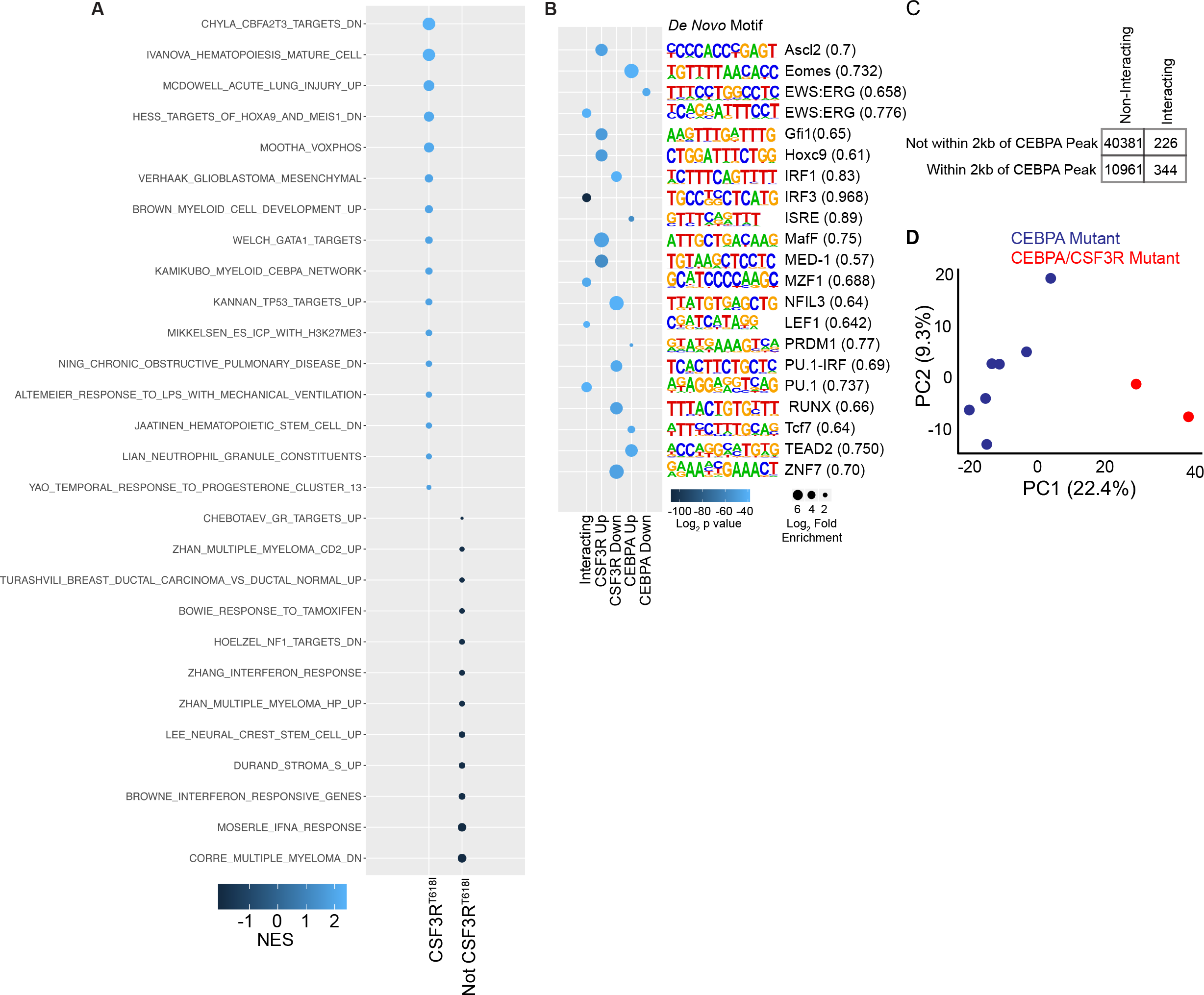
CEBPA mutations block myeloid differentiation. Related to Figure 2. **A.** Full GSEA results from Figure 2. All gene sets with q <0.05 shown. **B.** *De novo* motif enrichment at the promoters of differentially expressed genes from Figure 2 A. **C.** Permutation analysis of CEBPA ChIP-seq data and interacting gene subset, showing enrichment of CEBPA peaks within 2 kb of interacting genes (Fisher’s exact test, p<0.00001). **D.** PCA analysis of all genes differentially expressed between CEBPA mutant/CSF3R wild type and CEBPA mutant/CSF3R mutant pediatric AML.

**Figure S2.**
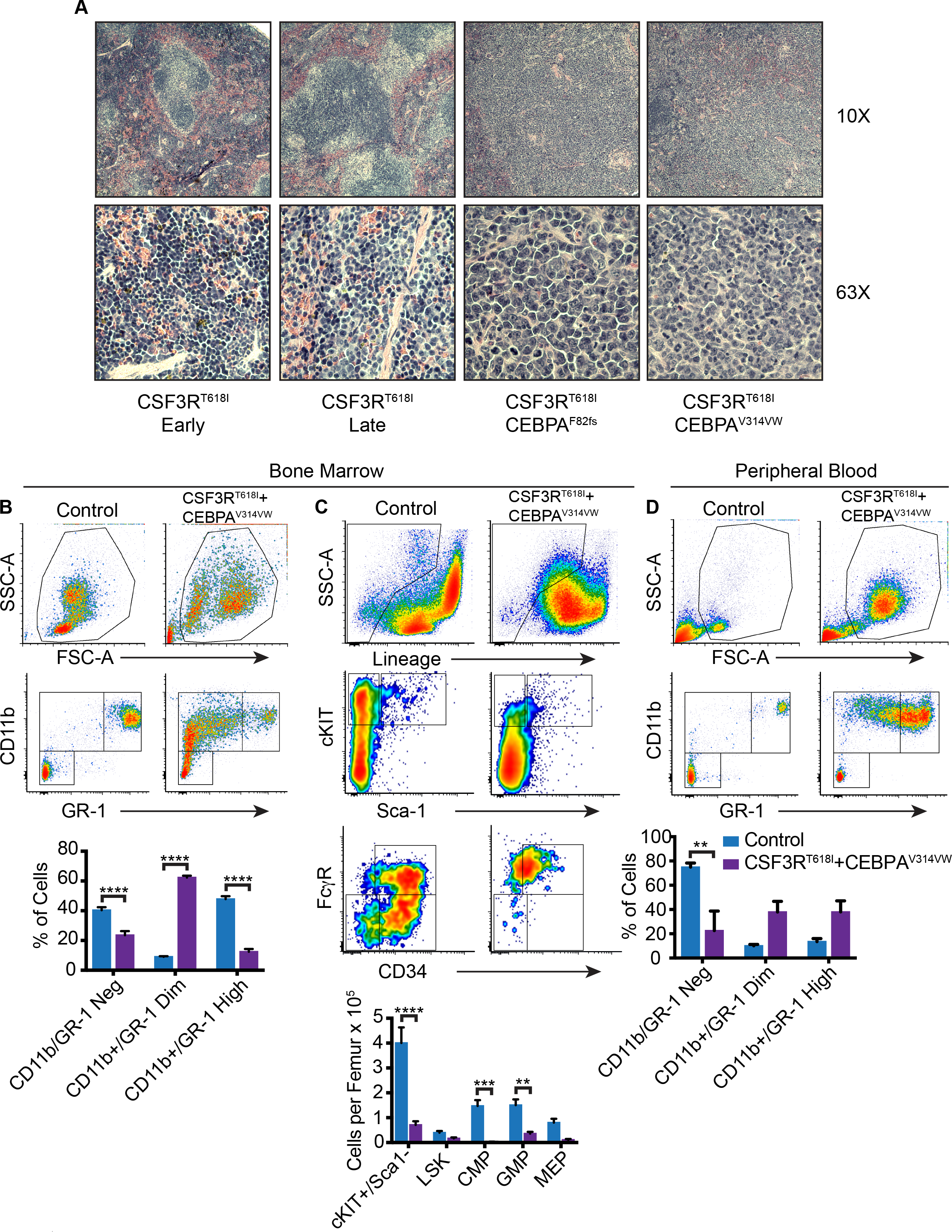
CSF3R^T618I^ and CEBPA^V314VW^ combine to produce an immature leukemia with replacement of normal hematopoiesis, while CEBPA^F82fs^ does not modify disease phenotype. Related to Figure 3. **A.** Representative images of spleens at low and high power from mice transplanted with CSF3R^T618I^ examined 3 weeks after transplant (Early) at time of overt disease development (~100 days, Late), CSF3R^T618I^ + CEBPA^F82fs^ at the time of disease development (120 days) and CSF3R^T618I^ + CEBPA^V314VW^ at the time of disease development (14 days post-transplant). Images representative of 3 animals per group (Same cohort described in Figure 2 A, B). **B.** Myeloid differentiation assessed by flow cytometry from bone marrow of BALB/c mice transplanted with 100,000 bone marrow cells transduced with CSF3R^T618I^ and CEBPA^V314VW^ at 14 days post-transplant as compared with control mice (Mice transplanted with empty vector are pancytopenic at day 14; n=5/group). **C**. Stem and progenitor cell populations from groups in B (n=5/group). **D.** Myeloid differentiation in the peripheral blood as assessed by flow cytometry for groups in B (n=4-5/group). In all cases, values are represented as mean +/− SEM. Statistical significance in all panels assessed by 2-way ANOVA with Sidak’s post-test, **: p<0.01, ***: p<0.001, ****: p<0.0001.

**Figure S3.**
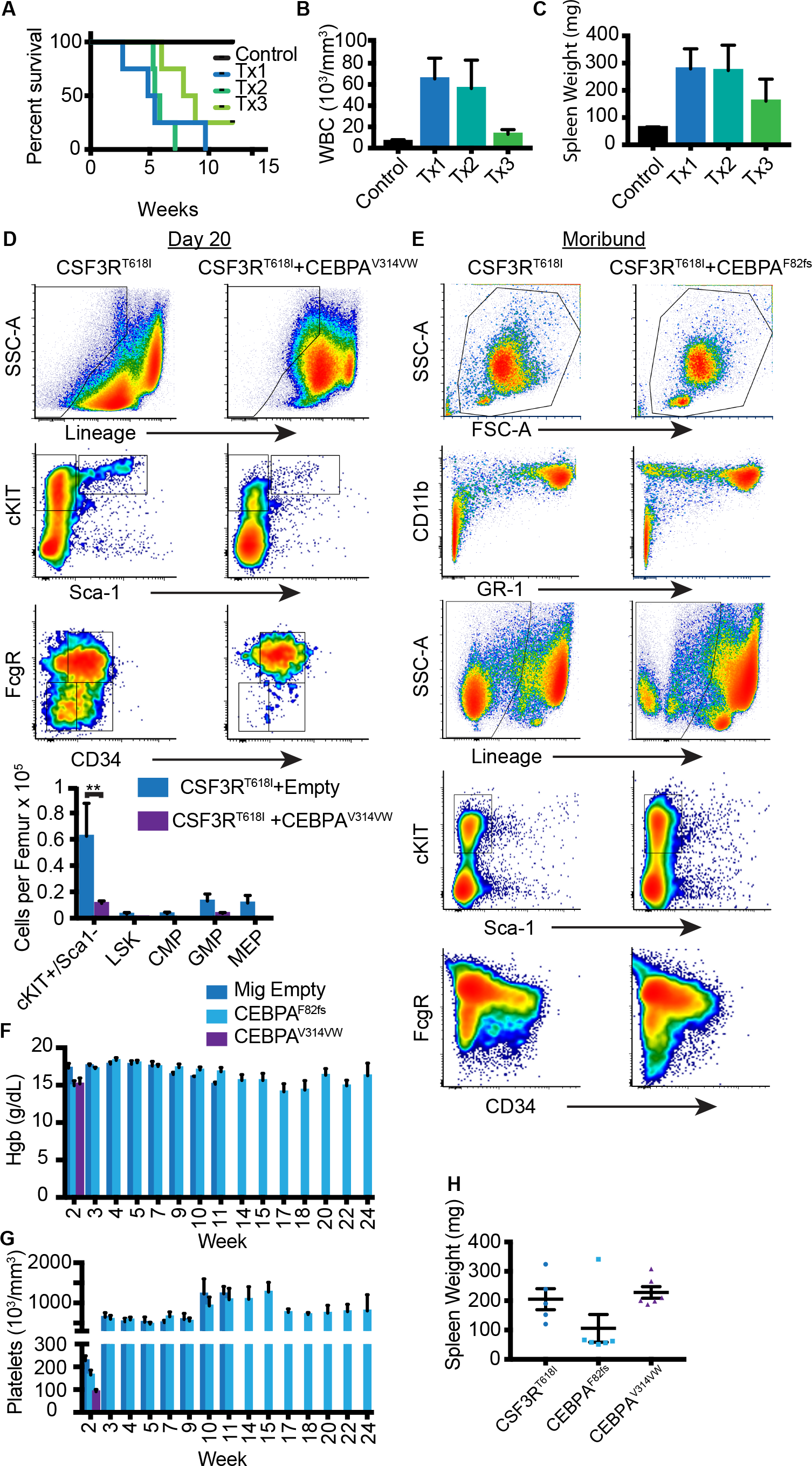
CEBPA^F82fs^ does not modify disease phenotype of CSF3R-dependent leukemia. Related to Figure 3. **A.** Survival after serial transplantation of 100,000 bone marrow cells from moribund primary recipients. Successive serial transplantation from moribund recipients for 3 serial transplantations (n=4-5/group). **B.** Terminal WBC counts from each round of serial transplantation. **C.** Spleen weight from each round of serial transplantation. **D.** Flow cytometric assessment of stem/progenitor cell populations at day 20 post-transplant in BALB/c mice transplanted with 10,000 cells transduced with CSF3R^T618I^ and CEBPA^V314VW^ compared with mice receiving 100,000 cells transduced with CSF3R^T618I^ and empty vector (At this time point, control mice had recovered from transplant-induced pancytopenia allowing direct comparison, same cohort as Figure 3C-G, n=3/group). **E.** Representative bone marrow flow cytometry for myeloid maturation and stem/progenitor cell populations from mice transplanted with CSF3R^T618I^ + Empty vector or CSF3R^T618I^ + CEBPA^F82fs^ at survival endpoint (Same cohort as Figure 3A, B, representative of 3 animals/group). **F.** Hemoglobin concentration over time in mice from Figure 2A, B. **G.** Platelet count over time in mice from Figure 3A, B. **H.** Spleen weight from mice in Figure 3A, B).

**Figure S4.**
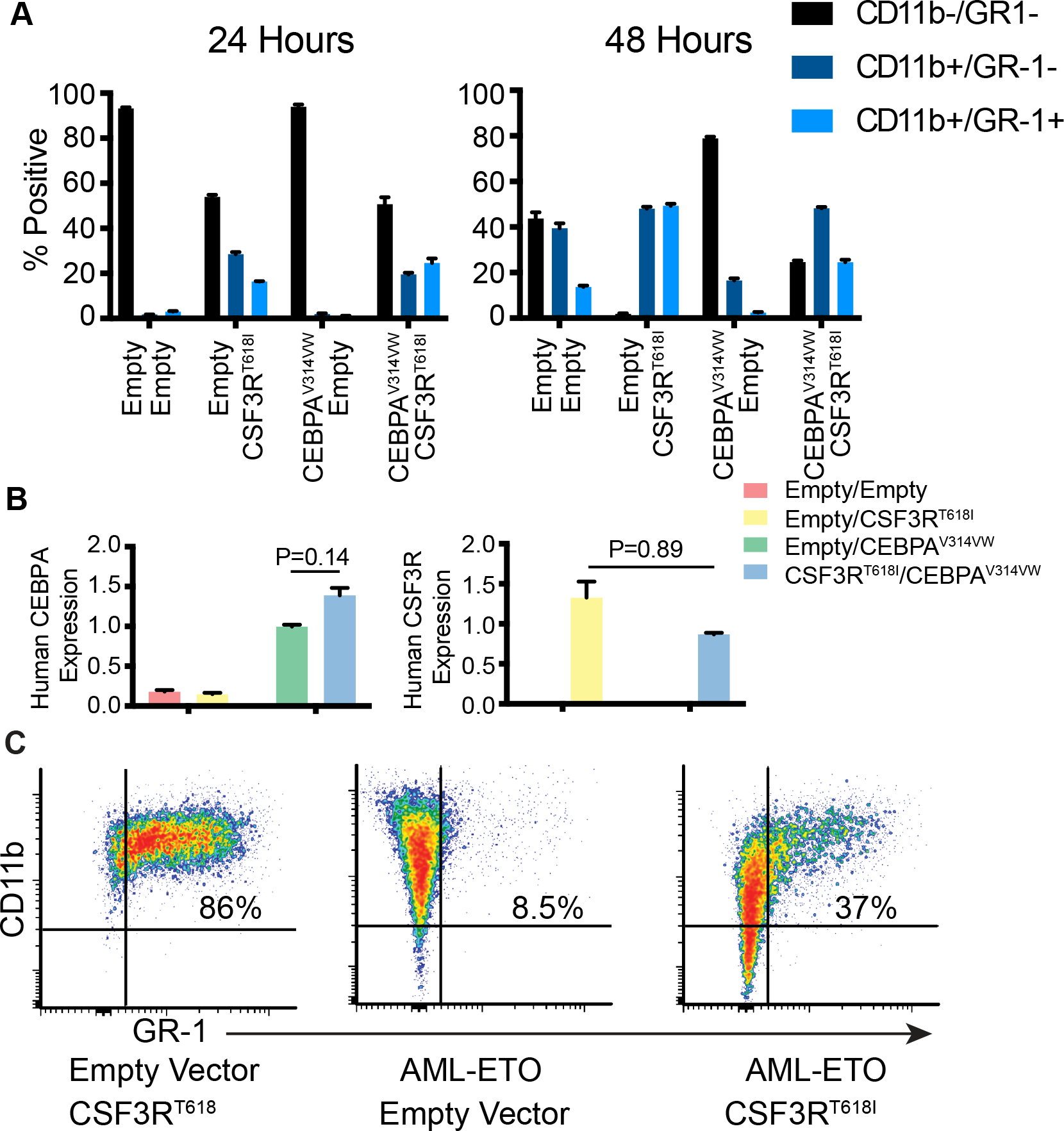
CSF3R^T618I^ and CEBPA^V314VW^ have opposing effects on myeloid differentiation in HoxB8 cells. Related to Figure 4. **A.** Quantification of percent of murine HoxB8-ER cells expressing GR-1 and CD11b as measured by flow cytometry after estrogen withdrawal. Cells were transduced with empty vector, CSF3R^T618I^, CEBPA^V314VW^ or both oncogenes in combination. **B.** Human CEBPA and CSF3R expression in HoxB8-ER cells from 3A, 24 hours after estrogen withdrawal as measured by Q-RT PCR. Statistical significance assessed by two-way ANOVA with Sidak’s post-test. (n=3/group). **C.** Expression of GR-1 and CD11b 48 hours after estrogen withdrawal as measured by flow cytometry in murine HoxB8 ER cells transduced with empty vector, CSF3R^T618I^ or AML-ETO. In all cases, values are represented as mean +/− SEM.

**Figure S5.**
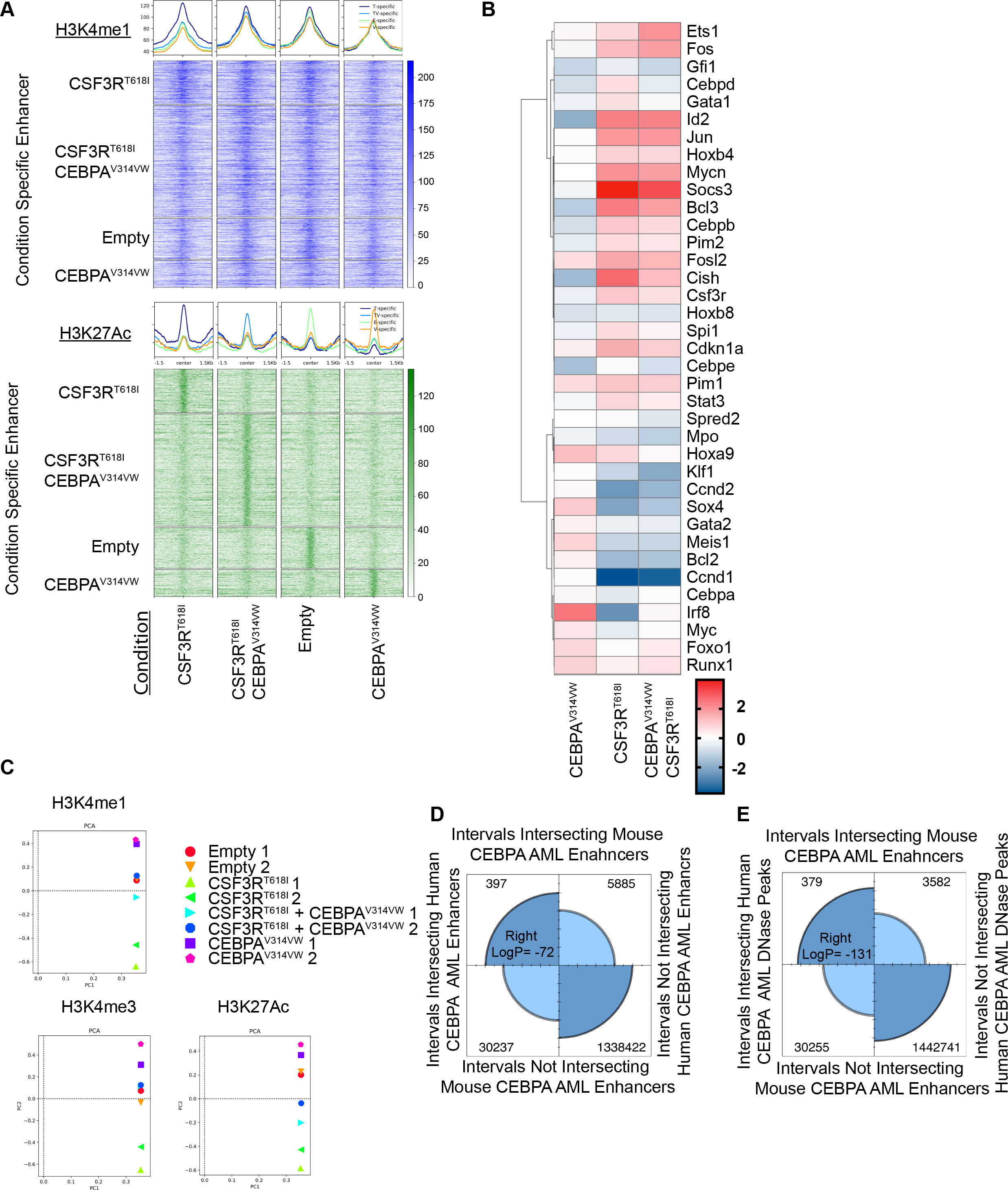
CEBPA Mutation Blocks Activation of CSF3R^T618I^ Specific Enhancers. Related to Figure 4. **A.** Heatmaps of H3K4me1 and H3K27Ac across each group of condition specific enhancers in all 4 treatment conditions**. B.** Confirmation of expression profile for specific genes identified in microarray analysis in Figure 5 E, performed by Taqman low density Q-PCR array, normalized to *GusB* and displayed as a relative quantity compared with empty vector control. All genes with differential expression (q<0.05) in at least one pairwise comparison shown. **C.** PCA analysis for H3K4me1, H3K4me3 and H3K27Ac across treatment conditions and experimental replicates. **D.** Overlap of enhancers identified in human CEBPA mutant AML with mouse CSF3R^T618I^/CEBPA^V314VW^ specific. **E.** Overlap of DNase I hypersensitivity peaks identified in Human CEBPA mutant AML with mouse CSF3R^T618I^/CEBPA^V314VW^ specific enhancers. D and E measured by Bedtools using Fishers-exact test. Right sided P value for significant enrichment listed.

**Figure S6.**
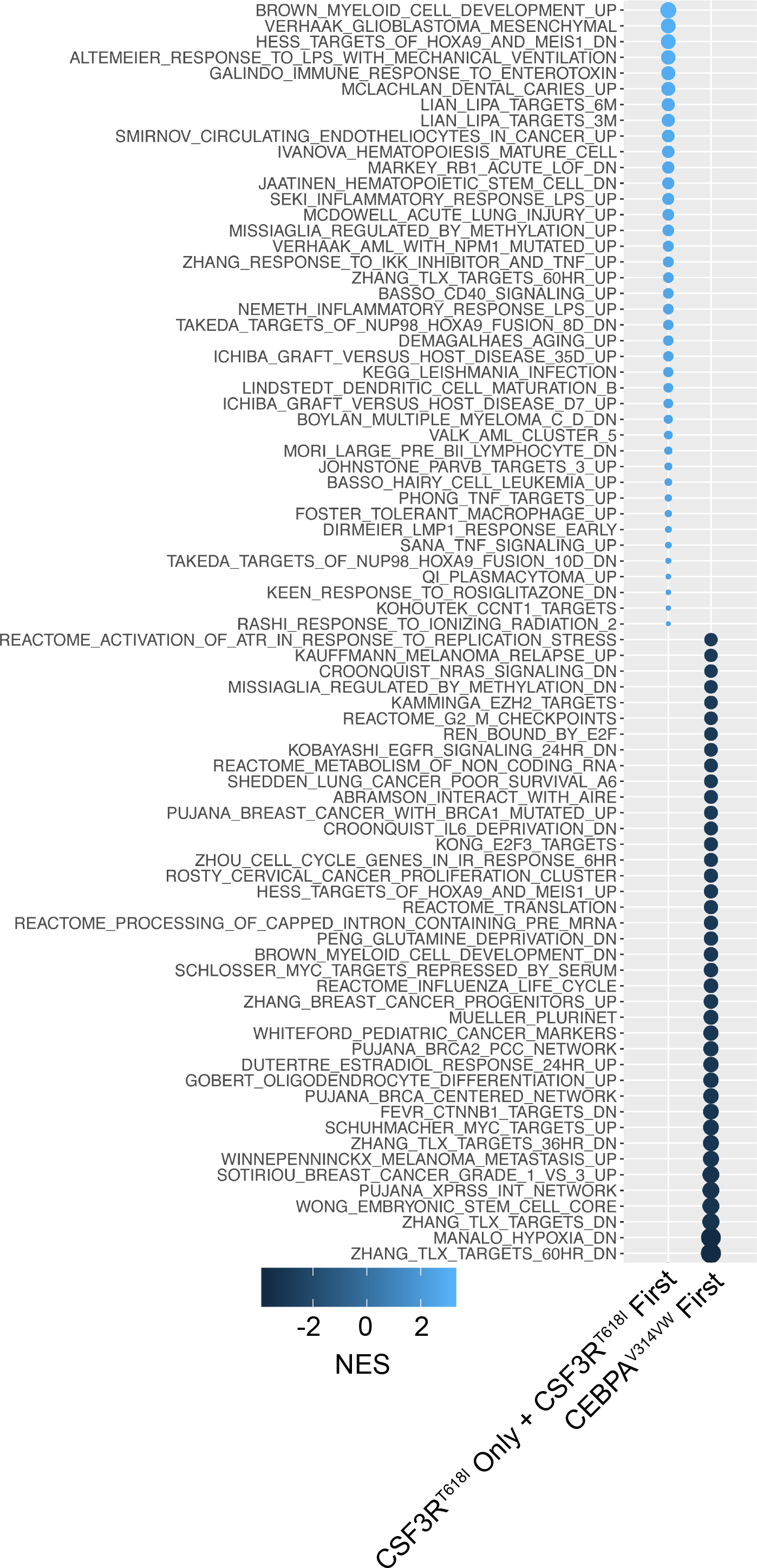
CEBPA First Blocks Myeloid Differentiation Gene Signatures Associated with CSF3R^T618I^. Related to Figure 5. Gene Set Enrichment Analysis results from RNA-seq data Figure 6 F. Top 40 C2 collection gene sets with q <0.05 enriched in CSF3R First/Only and enriched in CEBPA First.

**Figure S7.**
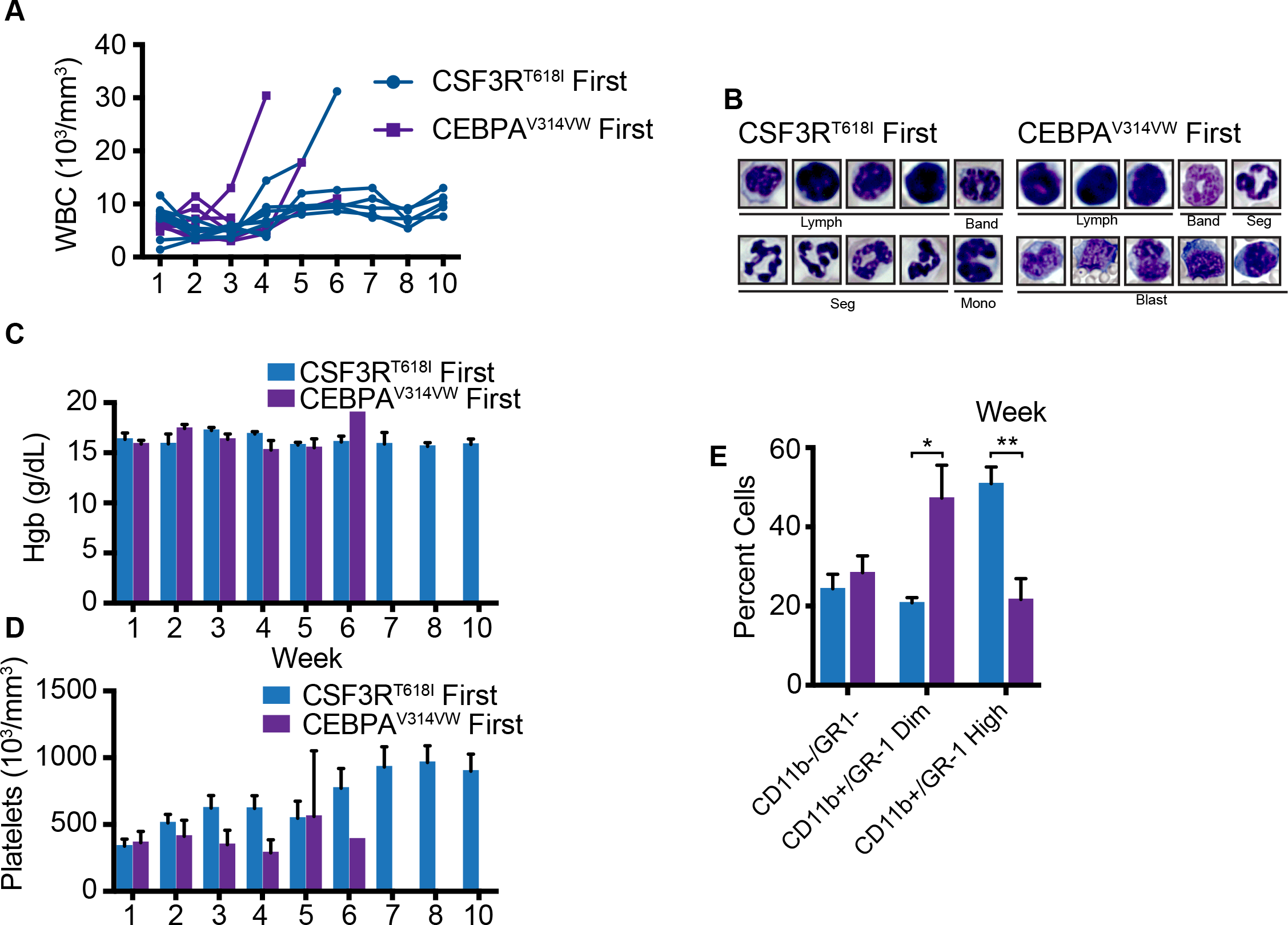
CEBPA^V314VW^ must precede CSF3R^T618I^ to produce a myeloid malignancy with differentiation block and cell cycle dysregulation. Related to Figure 6. **A.** WBC counts over time for mice in Figure 5. **B**. Representative images of cells for manual differential (representative of 3 animals/group). **C**. Hemoglobin over time in mice from Figure 5. **D.** Platelets over time in in mice from Figure 5. **E.** Quantification of flow cytometry from Figure 5 (n=3/group). In all cases, values are represented as mean +/− SEM. **: p<0.01 by Students t-test.

## METHODS

### Mice

C57BL/6J mice (#000664), Balb/cJ mice (#000651), Rosa26 ERT_2_-Cre mice (#008463) MX-1 Cre mice (#003556), and CEBPA^Flox/Flox^ mice (#006447) were obtained from The Jackson Laboratories. Female mice were used between 6-20 weeks, and age and weight matched in all experiments. Tamoxifen was dissolved in corn oil (Sigma) and injected intraperitoneally at 75 mg/kg/day for 5-days. Poly I:C (Sigma) was dissolved in PBS at 12 mg/kg on day 1, 3 and 5. All experiments were conducted in accordance with the National Institutes of Health Guide for the Care and Use of Laboratory Animals, and approved by the Institutional Animal Care and Use Committee of Oregon Health & Science University.

### Cell Lines

293T17 cells were obtained from ATCC and cultured in RPMI (Gibco) with 20% fetal calf serum (FCS, HyClone). Murine HoxB8-ER cells were a generous gift from David Sykes (Massachusetts General Hospital, Boston, MA) and cultured in RPMI (Gibco) with 10% FCS and CHO-SCF cell conditioned media (final concentration ~100 ng/mL) ^27^. Wild type HoxB8-ER cells were cultured and differentiated as previously described ^19^. Cell lines were tested monthly for mycoplasma contamination.

### Cloning and Retrovirus Production

The following plasmids were utilized: pMSCV-IRES GFP^7^, pMXs-IRES-Puro (Cell Biolabs Inc), pMSCV-IRES-mCherry FP (a gift from Dario Vignali, Addgene #52114), pMSCV-loxp-dsRed-loxp-eGFP-Puro-WPRE (a gift from Hans Clevers, Addgene #32702). The CSF3R^T618I^ mutation was generated previously ^7^. Full-length CEBPA and JAK3 cDNA was obtained from Genecopia. The CEBPA and JAK3 mutations were generated using the Quikchange Site Directed Mutagenesis Kit (Agilent) using the primers in Table S7. Mig containing AML-ETO was a gift from Dong-Er Zang, (Addgene #12434). To produce retrovirus, 293T17 cells were transfected with EcoPac helper plasmid (a gift from Dr. Rick Van Etten), and the appropriate transfer plasmid. Conditioned media was harvested 48-72 hours after transduction.

### Retroviral Transduction

Retroviral transduction of mouse bone marrow was performed as previously described ^7,29^. For colony assay, 2500 sorted cells were used per replicate. For bone marrow transplantation experiments, 200,000 cells were administered, with the balance of non-transduced cells as fresh syngeneic bone marrow. Survival endpoints for mutation comparison studies included WBC count>100, weight loss>20% initial weight, and moribund appearance.

### Real Time PCR

RNA was extracted using RNeasy or RNeasy micro kit (Qiagen). cDNA was synthesized using a High Capacity cDNA Synthesis kit (ThermoFisher). Real time PCR was performed using the QuantSudio7 Real Time PCR System (ThermoFisher) and Taqman primer probes (Thermofisher). For Taqman low-density arrays, array cards were custom printed by the manufacturer (ThermoFisher). Taqman primer probes used in this study are listed in Table S8.

### Flow Cytometry

The antibodies listed in Table S9 were utilized for FACS according to the manufacturer instructions. Stained cells were analyzed on a FACSAria III flow cytometer (BD).

### RNA Sequencing

Bone marrow from C57BL/6 mice was harvested and retrovirally transduced with the indicated oncogene combinations as described above. Cells were stained with a lineage cocktail, and lineage negative GFP/mCherry-double-positive cells were sorted. For order of acquisition experiments, cells were cultured in cytokine free I20 media for 48 hours after sorting in the presence of 500 nM 4-Hydroxytamoxifen. RNA was extracted using the RNeasy micro kit (Qiagen). cDNA libraries were constructed using the SMARTer universal low input RNA kit (Clonetech) and sequenced using a HiSeq 2500 Sequencer (Illumina) 100bp SR.

### Microarray

Labeled target, sscDNA was prepared from 12 RNA samples using the GeneChipTM WT-Plus protocol (ThermoFisher). Prior to cDNA synthesis and amplification, sample order was randomized. Amplified and labeled cDNA target samples were hybridized to a GeneChip Clariom S-Mouse array (ThermoFisher). Image processing was performed using Affymetrix Command Console (AGCC) v.3.1.1 software and expression analysis was performed using Affymetrix Expression Console. Differential Expression analysis was performed using the Transcript Analysis Console 4.0 (ThermoFisher).

### Chromatin-Immunoprecipitation Sequencing (ChIP-seq)

25 million cells per condition (Empty vector, CSF3R^T618I^-only, CEBPA^V314VW^-only, and CSF3R^T618I^ + CEBPA^V314VW^) were fixed with 1% formaldehyde and quenched by glycine. Cells were resuspended in lysis buffer (0.1% SDS, 0.5% Triton X-100, 20 mM Tris-HCl pH=8.0, 150 mM NaCl, 1x Proteinase inhibitor (Roche)), and sheared using the Bioruptor Pico sonicator (Diagenode). Reactions were rotated overnight at 4°C with antibody (H3K4me1 (ab8895, Abcam), H3K4me3 (ab8580, Abcam) andH3K27ac (ab4729, Abcam), Table S10). Next, samples were rotated with Protein A/G Magnetic Beads (Pierce). Beads were washed with TBST buffer (3x), lysis buffer, and 2xTE pH=8. Chromatin was eluted at room temperature in 1% SDS, 0.1 M NaHCO3. Crosslinks were reversed at 65°C overnight, digested with RNAse A and proteinase K, then purified by phenol-chloroform extraction. Sequencing libraries were generated using the NEBNext Ultra II DNA Library Prep Kit for Illumina (New England Biolabs). Libraries were sequenced using SE 75 bp Illumina NextSeq.

### Bioinformatic Analysis

#### RNA-seq Analysis: Murine Samples

Raw reads were trimmed with Trimmomatic^30^ and aligned with STAR^31^. Differential expression analysis was performed using DESeq2 ^32^. Raw p values were adjusted for multiple comparisons using the Benjamini-Hochberg method.

#### RNA seq Analysis: TARGET Pediatric AML Samples

RNA sequencing was performed on RNA collected from Pediatric AML samples as described in the initial TARGET AML publication^33^. Raw reads were aligned with Kallisto with count tables produced using Tximport^34,35^. Differential expression analysis was performed using DESeq2 ^32^. Raw p-values were adjusted for multiple comparisons using the Benjamini-Hochberg method. For comparison with mouse RNA seq, genes with differential expression between CSF3R WT and CSF3R^T618I^ were mapped to orthologous human genes using ensembl BioMart. Genes with differential expression in both mouse and human were considered and compared by Log2 Fold change between CSF3R^mutant^ and CSF3R^WT^ conditions.

#### Convergent Gene Expression Analysis: Leucegene AML

RPKM values from adult AML samples from the Leucegene cohort were downloaded using the following SRA accession numbers: GSE49642, GSE52656, GSE62190, GSE66917, GSE67039. Mouse genes with differential expression driven by CSF3R^T618I^ were converted to human gene symbols as above. Genes demonstrating an absolute fold change of >2 in both datasets were compared by Log2 Fold change between CSF3R^mutant^ and CSF3R^WT^ conditions.

#### Motif Enrichment Analysis

The enrichment analysis for motifs was performed using HOMER ^36^ using the - findMotifs command in a 500 bp window upstream and 200 bp downstream of the transcriptional start site. *De novo* motifs at enhancers were identified using the - findMotifsGenome command in HOMER in a +/− 1000 bp region surrounding the peak center. *De novo* motifs were matched to their closest known motif and displayed with the alignment score (with 1 being a perfect match). Top 5 motifs with p< 1E-10 for each group of promoters or enhancers are displayed.

#### Gene Set Enrichment Analysis

Gene set enrichment analysis using the GSEA software as previously described^37^. As CSF3R^T618I^-only and CSF3R^T618I^-first groups clustered together by Euclidian distance, they were combined and compared with CEBPA^V314VW^-first samples. Analysis was performed using the C2 collection from MSigDB. Permutations were performed by gene set and significance was set as an FDR adjusted p value of <0.05.

#### Permutation Analysis

CEBPA ChIP-seq peaks from GMPs ^15^ were converted from the mm9 to mm10, using the liftOver tool from the UCSC genome browser. We used BEDTools^38^ to randomly shuffle the location of all interacting genes, within their original chromosomes. After each shuffle, we used the closest-features subcommand in BEDOPS^39^ to calculate the distance between the new position of all shuffled genes and the closest CEBPA ChIP-seq.

#### ChIP-seq Analysis

Reads were aligned to the mouse reference genome (mm10) using bwa 0.7.12^40^ with default single end settings. Low mapping alignments were removed with samtools (MAPQ<30)^41^. Next, MACS2 2.1.1^42^ was used to predict significant peaks of ChIP-seq enrichment relative to the appropriate input controls and generate fold enrichment tracks. We used the ChromHMM software^43^ to characterize and annotate the genomes of each treatment group according to 6 chromatin states, based on different combinations of H3K4me1, H3K4me3 and H3K27ac marks. Enhancers were identified through the presence of H3K4me1 and H3K27Ac, absence of H3K4me3. Enhancers less than 500bp apart were merged. The closest gene to each enhancer was identified using BEDOPS^39^. In order to remove potential false positives due to proximity to large H3K27ac peaks, we removed treatment-specific putative enhancers without a H3K27ac peak summit.

#### Gene Ontology Analysis for ChIP-seq

Gene Ontology Analysis for histone mark ChIP-seq performed by Genomic Regions Enrichment of Annotations Tool ^44^ using the basal plus extension model to annotate enhancer coordinates with nearby genes.

#### Correlation of Gene Expression with Enhancer Activation

All differentially expressed genes as assessed by microarray analysis utilized for unsupervised hierarchical clustering analysis with a k-means=8. Each active enhancer was then annotated to the nearest gene. Condition-specific enhancers associated with each cluster of differentially expressed genes were counted and enrichment assessed as described under statistics.

#### H3K4me1 Differential Enrichment Analysis

To assess differences in levels of H3K4me1 across treatments, we used MACS2 bdgdiff subcommand^42^ to compare levels of H3K4me1 enrichment between the CSF3R^T618I^ and CEBPA^V314VW^ treatments. We set the minimum length of a region differentially enriched in H3K4me1 signal at 300 bp and merged differentially enriched regions less than 250 bp apart. Each region of differential H3K4me1 enrichment was annotated with its nearest active enhancer using BEDOPS^39^. Regions of differential H3K4me1 signal within 1kb of each group of condition-specific enhancers were counted. Enrichment assessed as described below under statistics.

#### Overlap of CEBPA Peaks with Condition-Specific Enhancers

We used publicly available genome-wide positions of all GMP CEBPA ChIP-seq peaks^15^. CEBPA peaks overlapping each group of condition-specific enhancers were identified using the mergePeaks command in HOMER with the -cobound option. CEBPA peaks associated with each group of condition-specific enhancers were counted and enrichment assessed as described below under statistics

#### Enhancer Overlap using BEDTools

Published human CEBPA AML DNase sensitivity peaks^26^ and enhancers^18^ were compared with mouse CSF3R/CEBPA-specific enhancers. The coordinates for mouse enhancers converted to human coordinates using the UCSC liftover tool. Overlap was assessed using the BEDTools fisher test^38^. Enrichment of overlap is detected using a fisher’s exact test with a significant right-sided p value while depletion is detected with a significant left-sided p value.

### Quantification and Statistical Analysis

Data are expressed as mean ± SEM. Statistical analysis was performed using Prism software (Version 7.0; Prism Software Corp.) or RStudio. Statistical analyses are described in figure legends. All data were analyzed with either an unpaired Student’s *t* test, or ANOVA followed by *post hoc* analysis using a Sidak’s corrected *t* test. For Taqman based array, microarray and RNA-seq data, p values were adjusted for repeated testing using a false discovery rate by the method of Benjamini-Hochberg ^45^. For enhancer enrichment analyses, a χ^2^ analysis was utilized with individual p-values adjusted by the method of Holm-Bonferroni. Survival analysis was conducted using the method of Kaplan–Meier and statistical significance was assessed using a Log Rank test.

### Data Availability

The accession numbers for all genomic data reported in this paper is GSE122166.

## SUPPLEMENTAL TABLES

**Table S1. RNA Sequencing Results for Simultaneous Oncogene Introduction.** Related to Figure 2.

**Table S2. RNA Sequencing Results for TARGET Pediatric AML Samples.** Related to Figure 2.

**Table S3. Microarray Results from Differentiating HoxB8-ER Cells.** Related to Figure 34

**Table S4. Activated Enhancer Catalog from Differentiating HoxB8 Cells.** Related to Figure 4.

**Table S5. Gene Ontology Analysis for Condition Specific Enhancers.** Related to Figure 4.

**Table S6. RNA Sequencing Analysis for Ordered Oncogene Introduction.** Related to Figure 5.

**Table S7.**
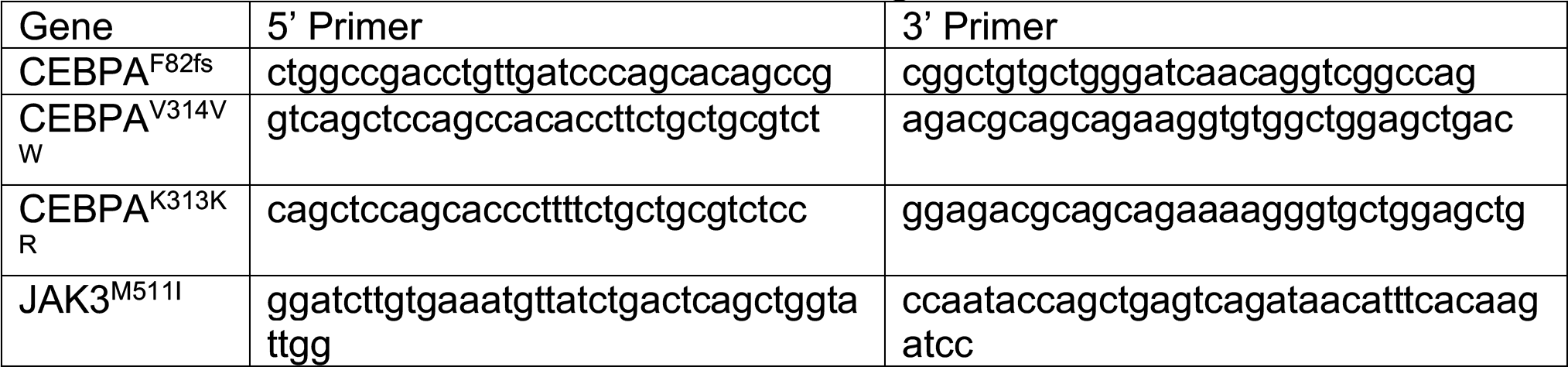
PCR Primers for Site Directed Mutagenesis.

**Table S8.**
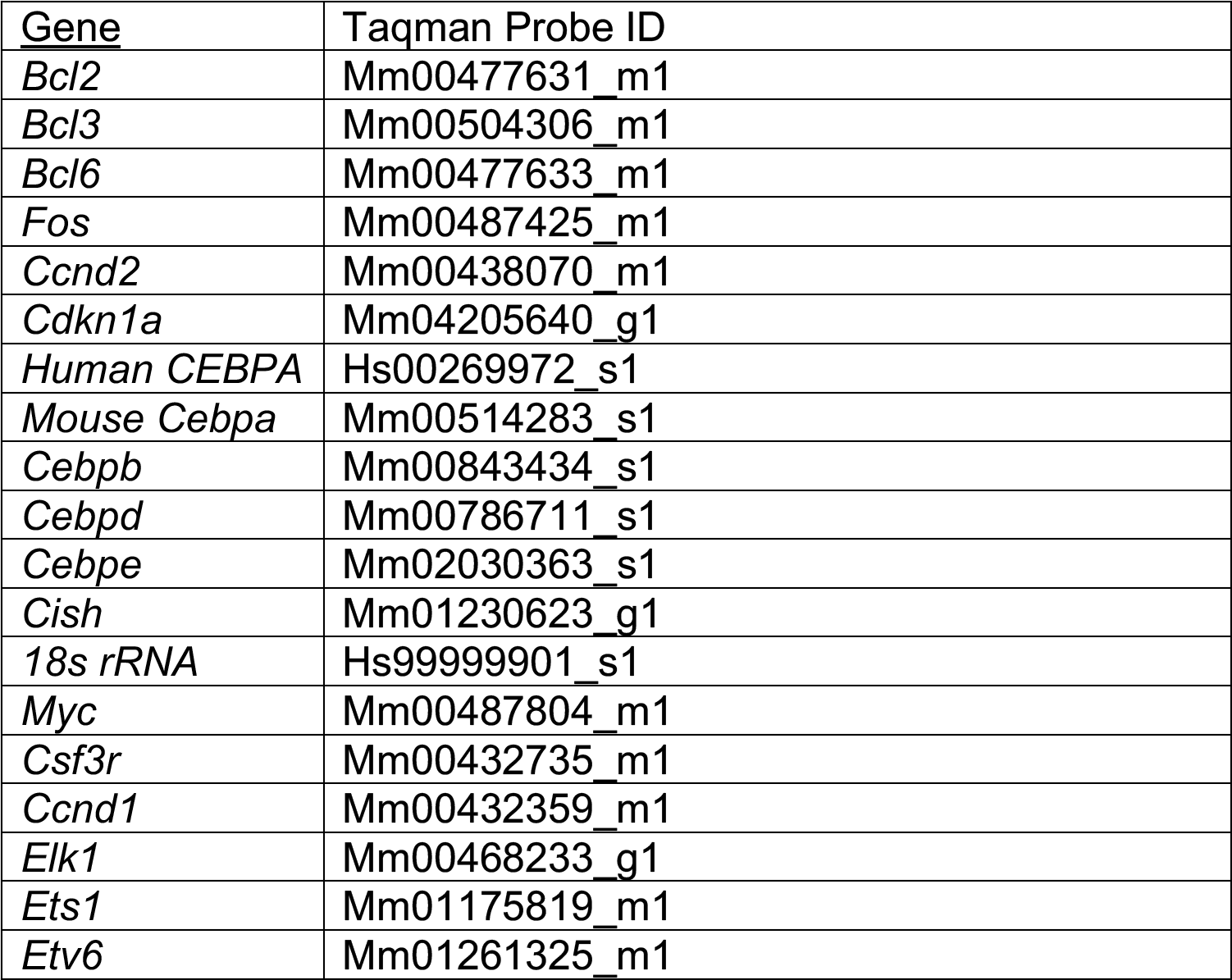
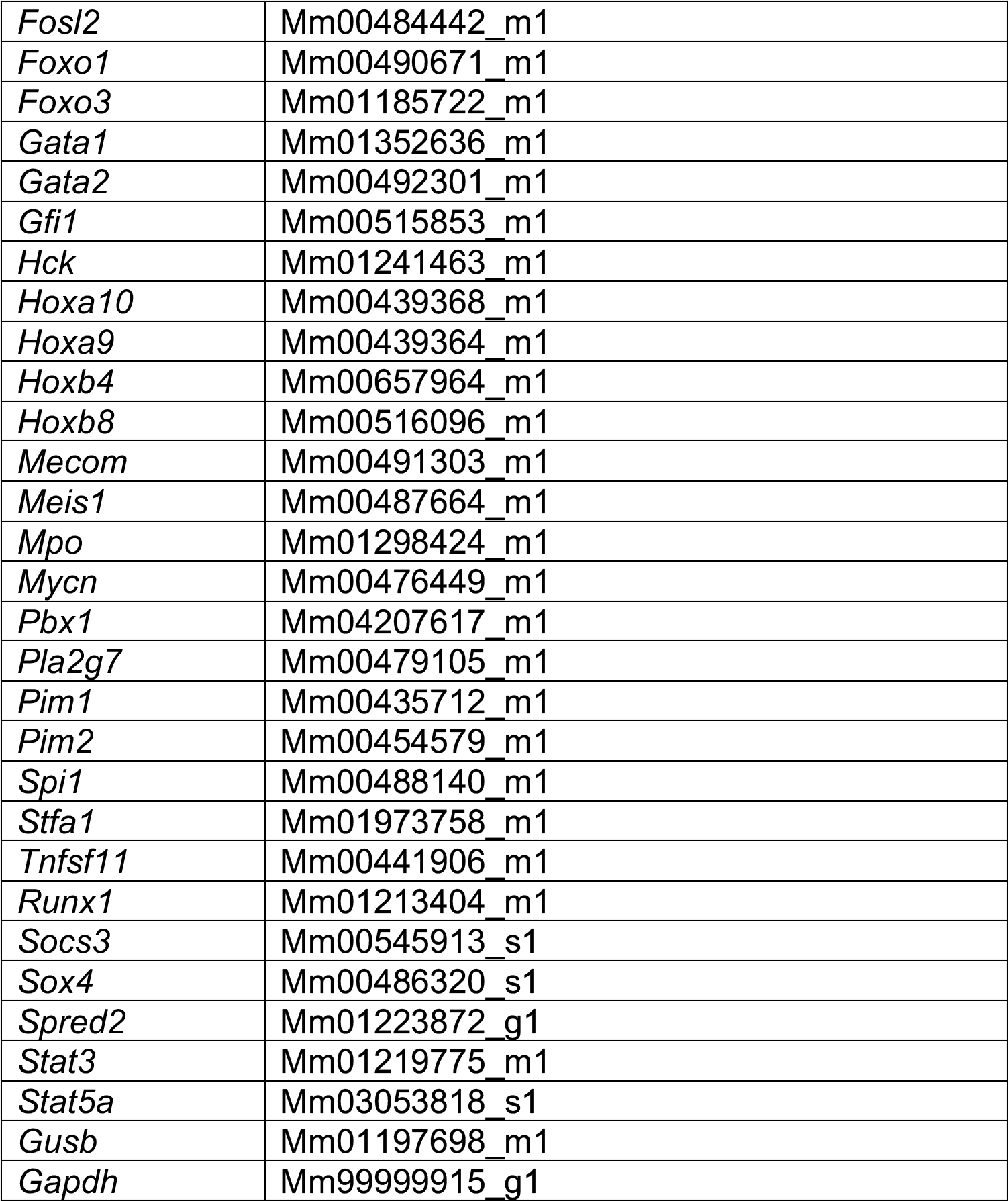
Taqman Primer Probes.

**Table S9.** Antibodies.

| Antibody | Manufacturer | Cat # | RRID |
| --- | --- | --- | --- |
| PE Rat Anti-Mouse GR-1 (RB6-8C5) | BD | 553128 | RRID:AB_394644 |
| PE-Cy7 Rat Anti-Mouse CD11b (M1/70) | BD | 561098 | RRID:AB_2033994 |
| PE-Cy7 Rat Anti-Mouse CD117 (2B8) | BD | 558163 | RRID:AB_647250 |
| PE Rat Anti Mouse Sca-1 (D7) | BD | 553108 | RRID:AB_394629 |
| APC Mouse Lineage Cocktail | BD | 558074 | RRID:AB_1645213 |
| BV421 Rat Anti-Mouse CD34 (RAM34) | BD | 562608 | RRID:AB_11154576 |
| PerCP-e710 Rat Anti-Mouse CD16/32 (93) | eBiosciences | 46-0161-80 | RRID:AB_2016632 |
| e450 Rat Anti-Mouse CD34 (RAM34) | eBiosciences | 48-0341-80 | RRID:AB_2043838 |
| Polyclonal Rabbit Anti-H3K4me1 | Abcam | ab8895 | RRID:AB_306847 |
| Polyclonal Rabbit Anti-H3K4me3 | Abcam | ab8580 | RRID:AB_306649 |
| Polyclonal Rabbit Anti-H3K27ac | Abcam | ab4729 | RRID:AB_2118291 |

